# Serine/Threonine phosphatase PP1 is a regulator of Notch signalling

**DOI:** 10.64898/2026.09.17.752308

**Authors:** Jiban Barman, Asif Bakshi, Rashmi Sipani, Rohit Joshi

## Abstract

Cell diversity generation is cardinal to the development of the functional central nervous system. The Notch pathway plays an important role in neurogenesis, spanning cell fate determination, cell death, and neural stem cell (NSC) competence switching, and is therefore highly regulated within cells. While the phosphorylation-based regulation of the pathway and its associated kinases is known, few phosphatases have been identified to counterbalance these regulations. Protein Phosphatase 1 is a member of a Serine/Threonine family of phosphatases responsible for a large majority of dephosphorylation events in the cell. In this study, we identify PP1-α and its regulatory subunit, PNUTS, as novel regulators of the Notch signalling pathway during *Drosophila* neurogenesis. We show that PP1-α87B/PNUTS positively regulate Notch signalling by dephosphorylating a highly conserved Serine residue in Su(H) to restore its DNA binding activity and thereby activating Notch downstream targets during neurogenesis. This facilitates the execution of two distinct physiological events, NSC apoptosis and competence switching, in different regions of the *Drosophila* CNS. We find that this regulation of the Notch pathway is also extendable to another cellular context, epithelial wing disc tissue, and critically relies on the phosphatase activity of PP1. Given that we can rescue the Notch-dependent depletion phenotypes of PP1-α using its human ortholog, we believe this regulation is likely to be conserved across species during development.

## Introduction

The Notch signalling pathway is an evolutionarily conserved, juxtacrine signalling pathway that plays a crucial role in binary cell-fate decisions during metazoan development (1). The pathway is also known to be involved in tissue homeostasis and generating cell diversity by regulating cell proliferation, differentiation, and apoptosis. It works through a membrane-localized Notch receptor (**N**) on the signal-receiving cell, which is activated by membrane-bound ligands, such as Delta/Serrate/Jagged (**Dl/Ser/Jag**). This activation leads to cleavage of the Notch intracellular domain (**NICD**), which then translocates into the nucleus and binds to the executive transcription factor (TF) **CSL** or **RBPJ** (C-promoter Binding Factor 1 in *H. Sapien*-**CBF1;** Suppressor of Hairless-**Su(H)** in *Drosophila*; Lin-12 and Glp-1-**Lag-1** in *C. elegans* or Recombination signal binding protein for immunoglobulin kappa J region-**RBPJ**) (1–4). NICD-CSL complex on DNA replaces corepressors (Hairy, Groucho, or CtBP) and recruits co-activator Mastermind-Like (**Mam/Maml**) and Histone AcetylTransferase (**p300/CBP**) to activate Notch-responsive downstream target genes like *Hes* family (Enhancer of Split complex [**E(spl)-C**] genes in *Drosophila*) (1–4).

Considering its importance in development, Notch signalling is tightly regulated at multiple levels (1, 5, 6), by a wide range of post-translational modifications of its components (7). Phosphorylation-dependent tuning of activation and repression of the Notch pathway is one such regulatory mechanism, which has been widely reported primarily for the Notch receptor and ligand (Dll1-Delta-Dl) (7) and, more recently, for the executive TF CSL (Su(H) in *Drosophila*) (8, 9). Depending on the residue, the phosphorylation of NICD can result in a reduction of NICD-Su(H)-Mam activation complex on the DNA (10) or affect NICD’s stability (7, 10–16). The phosphorylation of Su(H), on the other hand, has been shown to abrogate its DNA-binding ability on target DNA, thereby impacting signal transduction without affecting its stability (8, 9, 17). How the DNA binding of the phosphorylated Su(H) is restored to transduce the signal remains unknown. Most importantly, while various kinases have been shown to phosphorylate specific residues on different molecular players in the pathway (in a context-specific manner) (7–9), the relevant phosphatases that directly regulate the pathway components remain largely unknown. Eya1 is the only Phosphatase known to increase the stability of NICD-1 (7), whereas no such phosphatase has been reported for CSL.

Protein Phosphatase 1 (PP1) is a member of the Ser/Thr phosphoprotein phosphatases family, which includes other phosphatases such as PP2A, PP3 (PP2B), PP4, PP5, PP6, and PP7 (18). Amongst these, PP1 is a highly conserved, ubiquitously expressed oligomeric enzyme with catalytic and regulatory subunits (20, 21) and is well-characterised for its contributions to various cellular processes (19, 20). In vertebrates, the PP1 catalytic subunit has three canonical isoforms (α, β, and γ) encoded by different genes that exhibit a high degree of similarity and are predicted to generate multiple transcripts (21–23). *Drosophila* has orthologs of only the α and β catalytic subunits, which share high sequence identity but display isoform-specific functions due to differential regulatory interactions (24). The α catalytic subunit is coded by three genes (*PP1-α13C, PP1-α87B,* and *PP1-α96A*) (24), while the β catalytic subunit is coded by a single gene (*PP1-β9C* or *flapwing-flw*) (24, 25). The functional diversity and specificity of PP1 result from its tissue- and substrate-specific interactions and are controlled by a diverse set of regulatory subunits called Phosphatase-Interacting Proteins (PIPs). Compared to 200 PIPs known in vertebrates, *Drosophila* has only 40 PIPs (24, 26). One such PIP is PNUTS (**P**P-1 **NU**clear **T**argeting **S**ubunit) (27), which has been implicated in wide ranging PP1-dependent and independent roles. One of which is the recruitment of PP1 to different chromosomal loci, where it modifies RNA Pol II to regulate its transcriptional efficacy and control the expression of developmental genes (26, 28–30).

Bilaterian organisms require a complex CNS with region-specific cell diversity derived from a limited number of neural stem cells (NSCs); for this, NSCs need to integrate spatial and temporal information (31). The CNS of *Drosophila* comprise optic lobes (**OL**), central brain (**CB**), and ventral nerve cord (**VNC**) (32–34) (Fig. 1A). Within these regions, neurogenesis happens in two phases (embryonic and post-embryonic-larval/pupal) (35); during which Hox family of TFs partly regulate the spatial patterning of NSCs (or neuroblast-**NB**) (32–34), while the temporal information is provided by two parallel mechanisms: cascades of temporal transcription factors (**TTFs**) (32–34) and opposing temporal gradients of RNA-binding proteins (IGF-II mRNA-binding protein-**Imp** and Syncrip-**Syp**) (31, 35–41). In the early third instar larval (**eL3**) stage (60 hrs after larval hatching-ALH), NBs undergo a steroid hormone (Ecdysone) mediated temporal switching of their competence, following which they stop expressing Imp (marker for young NBs and TF Chinmo) and start expressing Syp (marker of old NBs and associated late TFs like Broad-Br and E93) (42) (Fig. 1A’). This temporal switching, also known as the Early to Late (**E>L**) competence switch, induces transcriptional changes in NBs, leading to their post-mitotic progeny differentiating into distinct neuronal subtypes. An equivalent developmental event marks an important step in vertebrate neurogenesis as well (43–49). Parallel to this, NBs in the abdominal and terminal segments (A3-A10) of larval VNC undergo Hox-dependent apoptosis which begins around mid L3 stage and by late L3 larval (**LL3**) stage all the NBs are eliminated (Fig. 1A). This developmental apoptosis regulates the number of neuronal progeny in these regions (50–58). (Fig. 1A). Here the resident Hox factor (Abdominal-A/Abdominal-B), helix-loop-helix TF Grainyhead (Grh) and Notch signalling pathway transcriptionally activate the *RHG* family of apoptotic genes through a 1Kb enhancer (*F3B3*) (54–57) (Fig. 1A). In *Drosophila* while the Notch signalling is required for apoptosis in most of the NBs, its role in E>L transition is reported to be restricted to 20% of these cells (59), and the molecular details of how Notch signalling executes the E>L transition in the NBs remains unknown.

**Fig 1.**
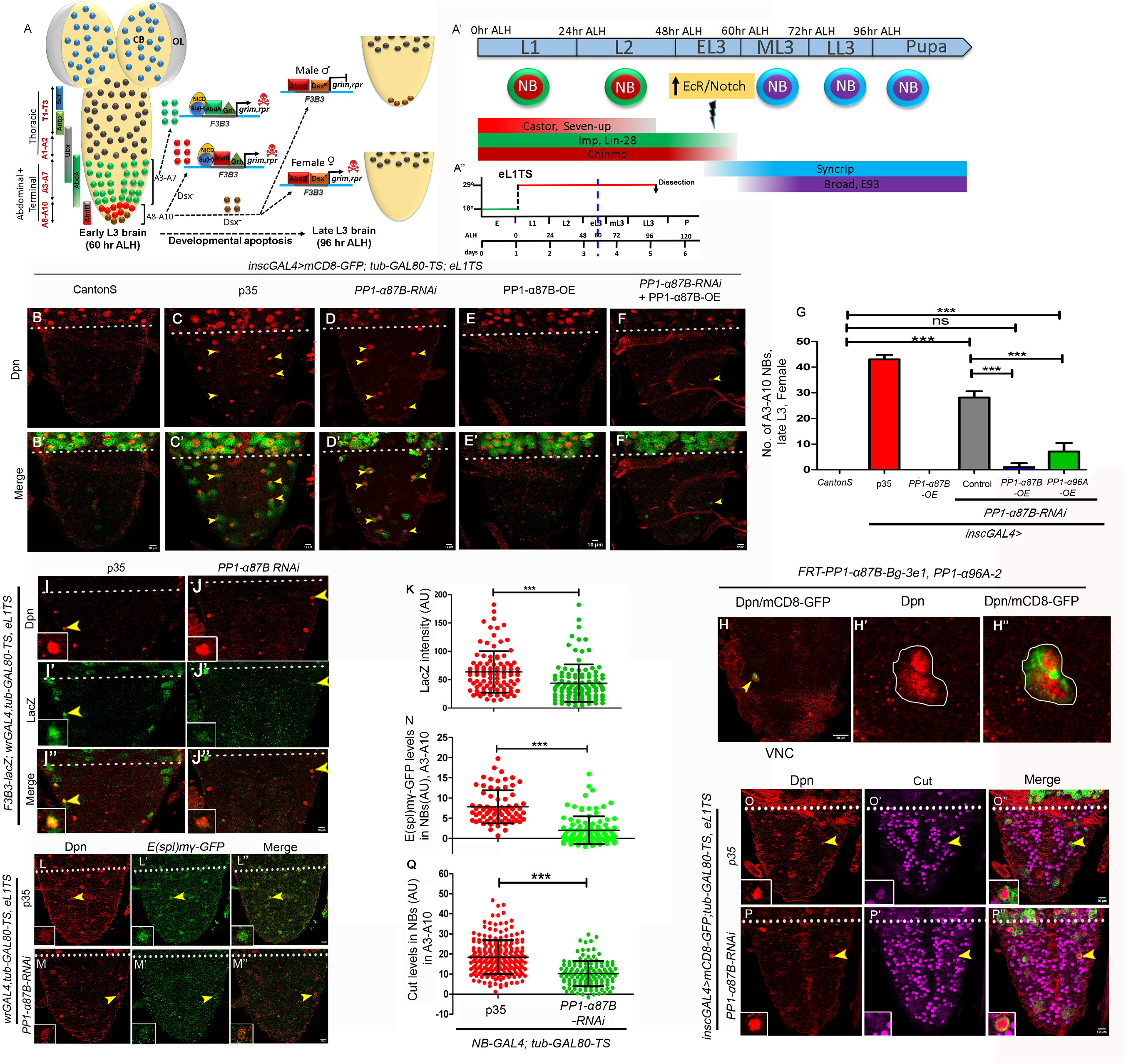
Protein Phosphatase 1α is a regulator of the Notch signalling pathway. (A) Schematic of larval CNS showing, showing optic lobe (OL), central brain (CB) and ventral nerve cord (VNC). Within the VNC segment-specific Hox-dependent NB apoptosis for 30 NBs in abdominal (A3-A7) and 12 NBs in terminal (A8-A10) segments is shown. In Abd-B expressing terminal (A8-A10) segments, NBs are categorised based on the expression of *doublesex* (*dsx*). In the female CNS, Dsx-positive NBs undergo apoptosis, whereas in males, these NBs continue dividing till late pupal stages. The Dsx-negative NBs of terminal segments (A8-A10 NBs) and abdominal NBs (in A3-A7 NBs) use Hox (Abd-B/AbdA), Grh and Notch signalling to transcriptionally activate the *RHG* family of apoptotic genes (*grim* and *reaper*) through a 1 kb enhancer (*F3B3*) and initiate apoptosis in early to mid L3 stages (60-72hrs after larval hatching-ALH). (A’) Schematic of larval NBs undergoing the temporal patterning programme. During early larval stages (till 56-60 hrs ALH), NBs express Early temporal factors (Castor, Seven-up, Imp, Chinmo, Lin-28). Subsequently, in response to an increase in levels of Ecdysone (Ec), EcR (Ecdysone Receptor) executes a competence switch in the NBs, leading to expression of Late temporal factors (Syncrip, Broad, E93, etc.). This switch is termed as the Early to Late (E>L) competence switch. A subset (∼20%) of the NBs in optic lobes use the Notch signalling pathway to execute the E>L switch (59). (A”) Shows the developmental time lines and early L1 (eL1) temperature shift (TS) protocols used in experiments shown in the figure. The L3 stage has been divided into two 16-hr intervals and a 24-hr interval to define the early, mid, and late L3 (eL3, mL3 and LL3) stages. The vertical blue dotted line indicates the approximate time of the E>L switch. (B-F) Shows the representative image of A3-A10 segments of the larval VNC of different genotypes at the LL3 stage. Block of NB apoptosis is marked by the presence of Dpn-positive cells, and rescue of the phenotype is scored by a reduction in Dpn-positive cells. (B) *Canton S* show no surviving NBs (0±0 NBs, n=7 VNCs, N=3). (C) Overexpression of *p35* (43.14±1.77 NBs, n=8 VNCs, N=3) and (D) knockdown of *PP1-α87B* (28.14±2.47 NBs, n=7 VNCs, N=3) result in block of NB apoptosis. (E) Overexpression of *PP1-α87B* does not block NB apoptosis (0±0 NBs, n=7 VNCs, N=3). (F) Overexpression of wild-type *PP1-α87B* rescues the block of NB apoptosis seen in the case of *PP1-α87B knockdown* (1.56 NBs, n=11 VNCs, N=3), establishing the specificity of the phenotype. (G) Graph showing the number of surviving NB in A3-A10 segments for the genotypes shown in panels B-F. (H) A 20X view of the A3-A10 segments of LL3 stage larval VNC showing a MARCM clone of a NB which is mutant for both *PP1-α87B* and *PP1-α96A* (*PP1-α87B^87Bg-3e1^, PP1-α96^A2^*) and fails to undergo apoptosis (n=3 clones, recovered from 3 VNCs). (H’ and H’’) Show the magnified view of the surviving NB. GFP marks the mutant clone, and NB is marked by Dpn. (I-J, L-M, O-P) Show that upon the knockdown of *PP1-α87B*, the levels of lacZ (I-J), *E(spl)mγ-GFP* (L-M), and Cut (O-P) are reduced in the surviving NBs when compared to p35 expressing controls blocked for NB apoptosis. This establishes that *PP1-α87B* knockdown blocks NB apoptosis by affecting the Notch signalling pathway, which executes apoptosis through activation of *F3B3* enhancer. (K, N, Q) Graph comparing the intensities of *F3B3-lacZ* (K)*, E(spl)mγ-GFP* (N) and Cut (Q) in A3-A10 NBs in *p35* expressing control VNCs versus VNCs with *PP1-α87B* knockdown. (K) *F3B3-lacZ* (63.77±36.65 AU, for 100 NBs, n=10 VNCs, N=3 vs 44.02±33.07 AU for 107 NBs, n=10 VNC, N=3), (N) *E(spl)mγ-GFP* (7.81±4.01 AU for 72 NBs, n=6 VNCs, N=3 vs 2.01±3.44 AU for 98 NBs, n=6 VNCs, N=3) (Q) Cut (18.39±8.45 AU for 220 NBs, n=8 VNCs, N=3 vs 10.22±6.30 AU for 132 NBs, n=10 VNCs, N=3). All representative images shown are single confocal sections from LL3-stage female VNCs. Insets show a magnified view of the NBs. Mean intensities are quantified in arbitrary units (AU). Scale bars are 10 µm, except H, which is 20 µm. Graph shows mean±SD. Significance (P-value) is from One-way ANOVA test and 2-tailed Student’s unpaired t-test. ALH means after larval hatching. Yellow arrowheads indicate NBs. The white dotted line separates thoracic (T) and abdominal (A) segments of the VNC.

In this study, we identify PP1-α/PNUTS as a novel modulator of the Notch signalling pathway required for executing NB apoptosis and E>L temporal switch during the larval phase of *Drosophila* neurogenesis. For NB apoptosis in A3-A10 segments of the VNC, we find that PNUTS is the regulatory subunit that helps PP1-α select and dephosphorylate the Notch executive TF Su(H) at a highly conserved Serine-269, thereby restoring its DNA-binding activity and activating its downstream target genes at least in two-thirds of these cells. This modulation of Notch signalling is extendable to the epithelial imaginal disc as well. In a different functional context of the competence switching (E>L transition) in thoracic NBs, we find that PP1-α is responsible for executing this transition in the majority (85%) of the NBs. Within this, a small subset of NBs (∼15%), which utilise the Notch signalling for this E>L switch, use PP1-α/PNUTS to regulate the pathway partly by dephosphorylating Su(H) at Serine 269 in 9% of the NBs (in the remaining 6% of the cells, PP1-α probably dephosphorylate an unidentified residue). We also demonstrate a context-specific deployment of E(spl)-complex members downstream of the Notch pathway to execute NB apoptosis and competence switching. Collectively, our results suggest that PP1-α/PNUTS regulates the Notch signalling pathway by dephosphorylating Su(H) in a context-dependent manner, thereby executing two physiologically distinct functions in NBs during larval neurogenesis. Considering that the block of NB apoptosis and the E>L transition arising from PP1-α depletion could be rescued by human PPP1, and relied on its phosphatase activity, we believe that this regulation of the Notch signalling pathway is likely to be conserved across species during development.

## Results

### Protein Phosphatase 1α is a regulator of the Notch signalling pathway

Normally, by the late third instar larval (LL3) stage, NBs in A3-A10 (abdominal: A3-A7 and terminal: A8-A10) segments of larval CNS undergo apoptosis, and no Dpn-positive cells (Dpn: a bHLH TF that marks NB) can be seen in these segments in females (Fig. 1A and 1B). We identified Protein Phosphatase 1α at 87B (*PP1-α87B*, CG5650) in an RNA interference (RNAi) screen for genes regulating NB apoptosis in A3-A10 segments of larval VNC. We tested three RNAi lines (BL32414, v35024, and BL67911) targeting different regions of the *PP1-α87B* cDNA and observed that two of these lines (BL32414 and v35024; detailed in the Methods) blocked NB apoptosis (Fig. 1D, BL32414 and Fig. S1A-B, v35024). For this, we used a temporally controlled UAS-GAL4-GAL80 system, where a temperature switch (TS) from 18 °C to 29 °C was used to induce the knockdown (here on referred to as TS-system/protocol) from the early L1 stage (early L1 Temperature Switch-eL1TS) and VNCs were analysed in the late LL3 stage (96 hrs ALH; indicated by black arrow in Fig. 1A’’). The expression of cell death blocker *p35* (a caspase inhibitor) was used to assess the total number of NBs undergoing apoptosis in these segments (Fig. 1C and bar 2, Fig. 1G). The p35-expressing A3-A10 NBs also served as a control in subsequent experiments in which the level of a reporter (or marker) had to be checked and compared for a gene knockdown or overexpression.

In *Drosophila*, the alpha (α) catalytic subunit of PP1 is coded by three very similar genes (*PP1-α13C, PP1-α87B,* and *PP1-α96A*) (24), which function redundantly in vivo (24). Therefore, we tested transcript levels of all three α-isoforms and observed a significant reduction in all them in *PP1-α87B* knockdown (BL32414) (Fig. S1G). This was also corroborated by the fact that single knockdowns of *PP1-α96A* and *PP1-α13C* also resulted in the block of NB apoptosis (Fig. S1C-S1F). Hereafter, all the experiments and analyses were conducted using the BL32414 line. In order to establish the specificity of the phenotype observed in the case of *PP1-α87B* knockdown, we rescued the block of NB apoptosis by individually overexpressing RNAi-resistant versions of *PP1-α87B* (TS protocol, Fig. 1A’’) (Fig. 1F and 1G) and *α96A* (Fig. 1G) (eL1TS, TS-protocol: Fig.1A’’). The rescue of the block of NB apoptosis in this and subsequent experiments was scored by a reduction in the number of Dpn-positive cells (NBs). We observed that overexpression of *PP1-α87B or PP1-α96A* resulted in a much smaller number of Dpn-positive NBs in the VNC (Fig. 1F and bars 5 and 6 in Fig. 1G), which corroborated the functional redundancy of the isoforms and ruled out off-target effects.

To further establish the role of PP1-α-isoforms in NB apoptosis, we performed a MARCM experiment using loss-of-function alleles of *PP1-α87B* (*PP1-α87B^Bg-3e1^*) and *PP1-α96A* (*PP1-α96A^2^*). The MARCM clones were induced at the eL1 stage (8 hrs ALH) and analysed at the LL3 stage (96 hrs ALH, detailed in Materials and Methods). We observed that single mutants did not show any block of apoptosis on their own, while the double mutant clones for the two α-isoforms (*PP1-α87B* and *PP1-α96A*) blocked NB apoptosis (Fig. 1H), suggesting that these two catalytic subunits functioned redundantly to execute NB cell death, and double knockout was required to block it.

The NB apoptosis in larval VNC relies on transcriptional activation of the *RHG* family of apoptotic genes (*grim* and *reaper*) through a 1Kb enhancer (*F3B3*) (Fig. 1A)(56). Therefore, we compared the expression of *F3B3-lacZ* in the p35 expressing control NBs with those resulting from the knockdown of *PP1-α* (TS-protocol: Fig.1A’’), and observed a significant reduction in lacZ intensity in *PP1-α* knockdown NBs (Fig. 1I-1K). Following this, we used a similar experiment to test whether the knockdown of *PP1-α* reduced the expression of the transcription factors (Grh and the resident Hox factor Abd-A/Abd-B) known to be critical for initiation of NB apoptosis in the abdominal and terminal segments of the VNC (54, 56). Here, we observed no significant reduction in the expression of Grh and Abd-B, and, on the contrary, observed an increase in expression of Abd-A (trigger for abdominal NB apoptosis) in the case of *PP1-α* knockdown (Fig. S1I and S1N). Similarly, we also examined the Notch signalling pathway using *E(spl)mγ-GFP*, which exhibits strong expression and is known to be responsive to Notch signalling in the NBs (60). In this case, we observed a significant reduction in *E(spl)mγ-GFP* levels, indicating that Notch signalling is compromised in cells with *PP1-α* knockdown (Fig. 1L-1N). This was further corroborated by a reduction in the expression of Cut (Ct: A Cut class of homeodomain-containing transcription factor), an established Notch signalling readout in epidermis and CNS (Fig. 1O-1Q) (61, 62). However, we did not observe any significant reduction in the levels of pathway components like NICD and Su(H) in the surviving NBs of the larval VNC (Fig. S2A-S2E). It is to be noted that the knockdown of *Notch* is known to cause a block of NB apoptosis (∼32 NBs in A3-A10 segments) (56)(54), similar to what is reported here in the case of *PP1-α87B* knockdown (28.14±2.47 NBs).

Collectively, our results established that PP1-α87B, PP1-α96A and PP1-α13C are required for executing NB apoptosis in larval VNC by modulating the Notch signalling pathway, most likely through enzymatic regulation of the pathway components. We also find that the three α-isoforms of PP1 function redundantly, and RNAi-mediated knockdown of *PP1-α87B* results in knockdown of all three PP1-α isoforms; therefore, PP1-α87B and PP1-α were used interchangeably hereafter.

### PP1-α regulates Notch signalling by modulating the phosphorylation state of Su(H)

Since PP1-α is a Ser/Thr Phosphatase, and we had not found any reduction in the expression of N and Su(H) in the NBs in the case of *PP1-α87B* knockdown (Fig. S2A-S2E), we considered the possibility that it might regulate the phosphorylation state of a protein in the Notch signalling pathway without affecting its expression levels. To this end, two recent studies have also demonstrated that phosphorylation of Thr-426 (T426) and Ser-269 (S269) regulates the DNA-binding activity of Su(H) (8, 17). Therefore, we hypothesised that PP1-α-mediated Su(H) dephosphorylation may regulate the Notch activity in NBs. A consensus RXXS/T motif has been proposed as a potential PP1-α dephosphorylation site (63). Interestingly, we found a single match for this putative dephosphorylation motif (residues 266-269-^266^RLRS^269^) (Fig. 2A) corresponding to an already reported Serine 269 phosphorylation in Su(H) (8, 9, 64). These studies had used Mass spectrometry (8) to show that Ser-269 is phosphorylated in vivo and the phospho-mimetic version of Su(H) abrogates Notch signalling during wing disc development and haematopoiesis (8, 9). Therefore, we tested the role of a phospho-mimetic version of Su(H) (UAS-Su(H)^S269D^) by overexpressing it from the embryonic stage and assessing its impact on apoptosis at the LL3 stage (TS protocol: Fig. 2P). We observed that the overexpression resulted in a partial block of NB apoptosis in A3-A10 segments (Fig. 2B and bar 2 in graph Fig. 2D’). We also observed a significant reduction in Notch signalling as assessed by the levels of *E(spl)mγ-GFP* (Fig. 2E-2G) and Cut (Fig. 2H and Fig. S2F-S2G) in the surviving NBs upon the overexpression of Su(H)^S269D^ (TS protocol: Fig. 2P’). The overexpression of the wild-type (UAS-Su(H)^WT^) or the phospho-dead (UAS-Su(H)^S269Q^) version of Su(H) did not block apoptosis (Fig. S2N). To further establish the role of Serine 269 in apoptosis, we used an allele where the endogenous Su(H) locus is replaced by an attP exchange with Su(H) mutated at Serine 269 to Aspartate (S269D) (64)(*attP-Su(H)^S269D^*). This phospho-mimic allele of Su(H) has also been shown to inhibit Notch signalling in wing discs and hematopoietic cells (64). We assessed the control heterozygous larvae of this attP allele (*Su(H)^S269D^*/+) at the LL3 stage (96 hrs ALH) and observed that they did not show any block of NB apoptosis in the A3-A10 segments of the VNC (Fig. 2D’). The age-matched homozygous larvae of the allele (*attP-Su(H)^S269D^*/*attP-Su(H)^S269D^*) showed a significant developmental delay and were found to be in the L3 stage at 96 hr ALH (as assessed by L3 spiracles, which had not everted out from the body wall) and died 16 hrs later. Therefore, we analysed them at 108 hrs ALH and observed a block of NB apoptosis (19.13±9.90 NBs) (Fig. 2D and 2D’), which was approximately two-thirds of the number of surviving NBs observed in the case of the *PP1-α87B* knockdown.

**Fig 2.**
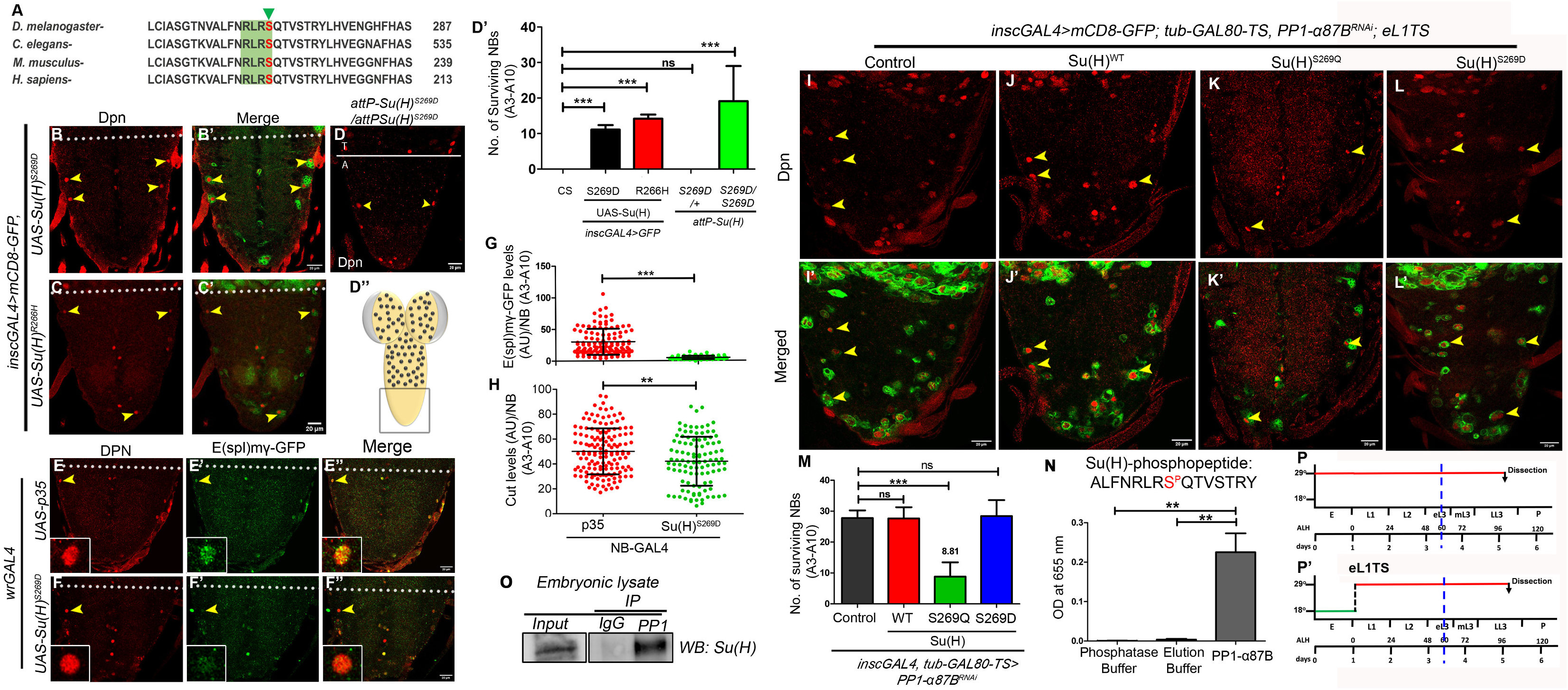
PP1-α regulates Notch signalling by regulating the phosphorylation state of Su(H). (A) Alignment of a region from *Drosophila* Su(H)-BTD (beta-trefoil domain) with its homologs from *C. elegans* (Lag1), *M. musculus* (RBPJ), and *H sapiens* (CBF-1). The conserved putative PP1-α phosphatase target site (R-X-X-S) is labelled with a green shadow highlighting the Serine 269 residue (*Drosophila*) in red. (B-C) Shows surviving NBs (Dpn-positive cells) in the A3-A10 segment upon overexpression of phospho-mimetic version (UAS-Su(H)^S269D^, 11.11±1.26 NBs, n=9 VNCs, N=3) and DNA-binding-deficient versions of Su(H) (UAS-Su(H)^R266H^, 14.20±1.13 NBs, n=10 VNCs, N=3) (TS: panel P). (D) Shows surviving NBs in larval brain homozygous for *attP-Su(H)* allele with Serine 269 replaced with an Aspartate at 108 hrs ALH (*attP-Su(H)^S269D^/attP-Su(H)^S269D^*:19.13±9.90 NBs, n=8 VNCs, N=4). (D’) Graph comparing the total number of surviving NBs in A3-A10 segments, observed in genotypes shown in panels B-D. (D’’) Region of the CNS shown in all the panels of the figure is indicated by the box. (E-F) Show that upon the overexpression of *Su(H)^S269D^*, the levels of *E(spl)mγ-GFP* are reduced in the surviving NBs in A3-A10 segments, when compared to p35 expressing controls (TS: panel P’). This establishes that overexpression of *Su(H)^S269D^* blocks NB apoptosis by altering Notch signalling. (G-H) Graph comparing the levels of *E(spl)mγ-GFP* (G) and Cut (H) in A3-A10 NBs overexpressing Su(H)^S269D^ with *p35* expressing control. *E(spl)mγ-GFP* (*p35*: 30.30 ± 1.96 AU for 116 NBs, n=12 VNCs, N=3; *vs Su(H)^S269D^*: 4.743± 0.34 AU for 62 NBs n=11 VNCs, N=3) and Cut (*p35*: 50.11±18.49 AU for 149 NBs, n=10 VNCs, N=3 vs *Su(H)^S269D^*: 42.19±19.65 AU for 108 NBs, n=13 VNCs, N=3). (I-L) Shows that the overexpression of the phospho-deficient version of Su(H) (*Su(H)^S269Q^*: 8.81±4.62 NBs, n=12 VNCs, N=3) (K) could partially rescue the block of NB apoptosis reported in case of *PP1-α87B* knockdown phenotype (27.80+-2.4, n=10 VNC and N=3) (I), but overexpression of wild type (*Su(H)^WT^*: 27.64±3.64 NBs, n=11 VNCs, N=3) (J), and constitutively phosphorylated versions of Su(H) (*Su(H)^S269D^* :28.42 ± 5.17 NBs, n=11 VNCs, N=3) (L) failed to do so. This suggests that Su(H) is downstream of PP1-α. (M) Graph showing the number of surviving NBs (Dpn-positive cells) in A3-A10 segments for the genotypes shown in panels I-L. (N) Graph showing the Malachite Green-based quantification of inorganic phosphate released from the Su(H) phosphopeptide (*D. melanogaster:* ALFNRLRS^P^QTVSTRY) by bacterially purified PP1-α87B in an in-vitro phosphatase assay (n=3). This establishes that PP1-α87B can dephosphorylate Su(H) at Serine 269. (O) Western blot showing that antisera for PP1-α87B could pull down Su(H) from embryonic lysate while IgG could not, suggesting that PP1-α87B and Su(H) exist as a complex in vivo. Gels and blots shown are representative of 3 repeats. (P-P’) Show the temperature shift (TS) protocols used for overexpression experiments shown in panels B-C, E-F (P) and panels I-L (P’). All representative images shown are single confocal sections from LL3-stage female VNCs. Mean intensities are quantified in arbitrary units (AU). Scale bars are 20 µm. Graph shows mean±SD. Significance (P-value) is from the One-way ANOVA test and 2-tailed Student’s unpaired t-test. ALH means after larval hatching. Yellow arrowheads indicate NBs. The white dotted line separates thoracic (T) and abdominal (A) segments of the VNC.

Since the phospho-mimetic version (Su(H)^S269D^) loses its DNA binding activity in vitro (8), we wanted to establish that the DNA binding activity of Su(H) was indeed required for NB apoptosis. To this end, we used a version of Su(H) which is incapable of binding to DNA in vitro (Su(H)^R266H^) (65). The overexpression of (UAS-Su(H)^R266H^) from embryonic to LL3 stage (TS protocol: Fig. 2P) also resulted in a partial block of NB apoptosis (Fig 2C and bar 3 in graph Fig. 2D’). We also observed a significant reduction in Notch signalling as assessed by levels of *E(spl)mγ-GFP* and Cut in the surviving NBs when compared to p35 expressing control NBs (Fig S2M and S2M’) (TS protocol: Fig. 2P’).

Next, we probed the hierarchy between Su(H) and PP1-α87B by overexpressing wild-type [UAS-Su(H)^WT^] and different phospho-state versions of Su(H) [UAS-Su(H)^S269D^ and Su(H)^S269Q^] in the background of *PP1-α87B* knockdown, and assessed their capacity to rescue the block of NB apoptosis caused by *PP1-α87B* knockdown (TS protocol: Fig 2P’). We observed that while the overexpression of Su(H)^WT^ (Fig. 2J and 2M) and constitutively phosphorylated version [Su(H)^S269D^] (Fig. 2L and 2M) did not rescue the NB apoptosis block observed in the case of *PP1-α87B* knockdown (Fig. 2I and 2M), the phospho-deficient version [Su(H)^S269Q^] could largely rescue it (Fig. 2K and 2M). This suggested that Su(H) is downstream of PP1-α87B. The experiment also suggests that normally Su(H) is phosphorylated at Serine 269 by a kinase. Following this phosphorylation, Su(H) dissociates from the DNA and needs to be dephosphorylated by PP1-α to regain DNA binding and to resume Notch signalling. Hence, WT and S269D versions are unable to rescue *PP1-α87B* knockdown, as Su(H) (in both cases) is likely to be locked in a phosphorylated state and unable to associate with DNA. On the other hand, S269Q cannot be phosphorylated and therefore may not fall off the DNA as easily in the first place; and as a result, Notch signalling is active, and hence this version alone can rescue the block of apoptosis caused by *PP1-87B* knockdown.

To conclusively establish that Su(H) is a bona fide target of PP1-α, we did an in vitro Phosphatase assay. For this, we used a custom-synthesized phosphopeptide of Su(H) (^262^ALFNRLRS^P^QTVSTRY^276^) with serine phosphorylation at position 269 (8, 9, 17). Here, we observed that bacterially purified PP1-α87B could release inorganic phosphate from Su(H) specific phosphopeptide (quantified using Malachite green at 655nm), establishing that PP1-α87B is capable of efficiently dephosphorylating Ser-269 of Su(H) (Fig. 2N).

Since PP1-α87B dephosphorylates Su(H), next, we expected that they may form a complex in vivo. For this, we used PP1-α87B as bait and successfully pulled down Su(H) from whole embryonic lysate using an anti-PP1-α87B antibody (Fig. 2O).

Collectively, our results from the rescue experiment with phospho-mimetic and phospho-dead versions of Su(H) establish that the Notch pathway is downstream of PP1-α and the DNA-binding activity of Su(H) is important for its ability to activate Notch targets and execute NB apoptosis. Our in vitro phosphatase assay and pulldown experiments further support the notion that PP1-α binds to Su(H) and modulates its DNA-binding activity by dephosphorylation at Ser-269.

### E(spl)-complex consolidates Notch response for execution of NB apoptosis

E(spl)-complex genes are crucial for neurogenesis and are the earliest targets expressed in response to Notch-signalling (66–68). The complex has 7 HLH TFs (mβ, mγ, mẟ, m3, m5, m7, and m8) and 4 Bearded family of transcriptional repressors (mα, m2, m4, and m6) (Fig. 3A). Amongst the HLH members, m8, mβ, and mγ are known to be expressed in *Drosophila* NBs (60, 69, 70). Since Notch signalling is required for NB apoptosis in VNC (56), we checked the expression of the remaining E(spl)-HLH members (mẟ, m3, m5, m7), using endogenously tagged reporter lines in A8-A10 NBs (70). We observed that only m3 showed expression in NBs in the mid L3 stage (in addition to m8, mβ, and mγ) (Fig. S3A-S3D). We also observed that since NBs initiate apoptosis asynchronously (over 48 hours from early to late L3), not all the NBs expressed the reporters simultaneously. Following this, we tested the role of the complex in NB apoptosis using MARCM. We could recover Dpn and GFP-positive homozygous mutant clones for *E(spl)-*C deletion (*E(spl)-C^Df^*(*^3^*)*^32.2 gro+^*) (71) in A3-A10 segments (Fig. 3B and 3E’’), establishing its role in NB apoptosis.

**Fig 3.**
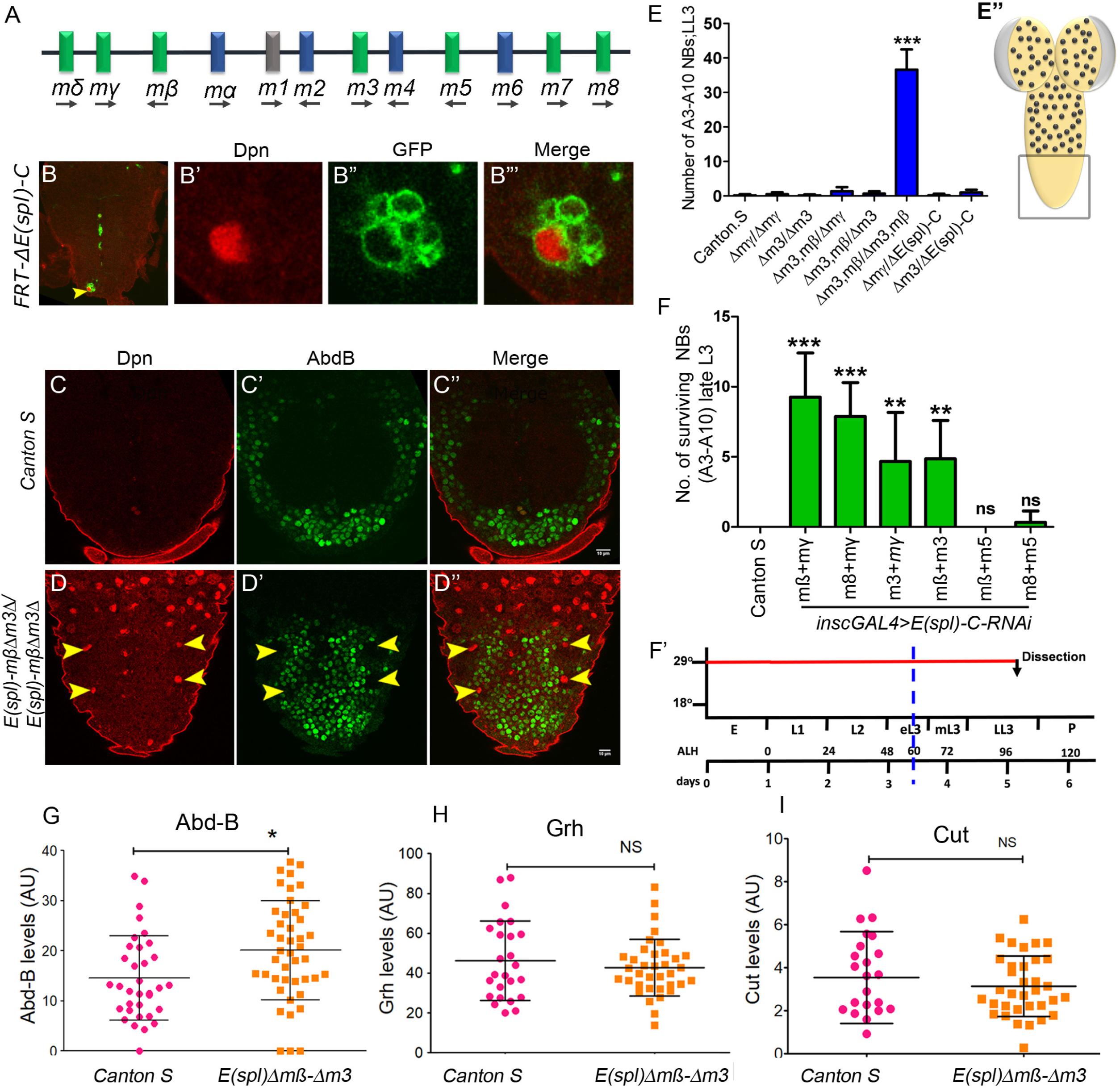
E(spl) complex members mß, m3 function downstream of the Notch pathway for execution of NB apoptosis. (A) Schematic shows the organisation of E(spl)-complex with bHLH family (Green) and Beard family (Blue) members in the (Ch-3R). The line represents genomic DNA, and the arrows indicate the direction of the transcripts. (B-B’”) MARCM clone of NB homozygous for *E(spl)-C* deletion does not undergo apoptosis and could be visualised in A3-A10 segments of LL3 stage VNC (n=14 clones, recovered from 8 VNCs). (B) 20X view of the A3-A10 segment of larval VNC. (B’-B’’’) Show the magnified view of the NBs. GFP marks the mutant clone, and Dpn marks NB. (C-D) Shows A3-A10 segments of LL3 stage VNC from *CantonS* controls with no surviving NBs (0.14±0.37 NBs, n=7 VNCs, N=2) while the age-matched VNCs (of LL3 stage) homozygous for double deletion of *E(spl)-Δmß,m3* (*Δmβm3/Δmßm3*) (36.60±5.87 NBs,n=10 VNCs, N=3) showed a block of NB apoptosis. (E) Graph showing the number of surviving NB in A3-A10 segments for control (*CantonS*) and the deletion combination analysed for the HLH-members of *E(spl)-C*. (E’) Region of the CNS shown in all the panels of the figure is indicated by the box. (F) Graph comparing the total number of surviving NBs in A3-A10 segments, observed in wildtype (*Canton S*: 0±0 NBs, n=8 VNCs, N=2) and double knockdown done for various members of E(spl)-C (*mβRNAi+mγRNAi*: 9.25±1.12 NBs, n=8 VNCs, N=3; *m8RNAi+mγRNAi*: 7.87±2.41 NBs, n=8 VNCs, N=3; *m3RNAi+mγRNAi*: 4.66±3.50 NBs, n=6 VNCs, N=2; *mβRNAi+m3RNAi*: 4.85±2.73 NBs, n=7 VNCs, N=2; *mβRNAi+m5RNAi*: 0±0 NBs, n=9 VNCs, N=3; *m8RNAi+m5RNAi*: 0.33±0.81 NBs, n=6 VNCs, N=2). (F’) Show the temperature protocols used for the double knockdown experiment of E(spl)-C complex shown in panel F. (G-I) Graph comparing the intensities of Abd-B (G), Grh (H) and Cut (I) in A8-A10 NBs in VNCs of *CantonS* versus double mutant homozygotes of *E(spl)-Δmß,Δm3* in mid L3 stage NBs. (G) Abd-B (*CantonS*: 20.11±9.9 AU, 34 NBs, n=5 VNCs, N=2 vs *E(spl)-Δmß,Δm3:* 14.58±8.45 AU, 41 NBs, n=5 VNC, N=2), (H) Grh (*CantonS:* 47.26±19.9 AU, 23 NBs, n=5 VNCs, N=2 vs *E(spl)-Δmß,Δm3:* 43.56±13.75 AU, 34 NBs, n=5 VNCs, N=2) and (I) Cut (*CantonS:* 3.52±2.13 AU, 23 NBs, n=5 VNCs, N=2 vs *E(spl)-Δmß,Δm3:* 3.14±1.32 AU, 34 NBs, n=5 VNC, N=2). The experiment suggests that Grh, AbdA and AbdB levels are unchanged in homozygotes of *E(spl)-Δmß,Δm3.* All representative images shown are single confocal sections from LL3-stage female VNCs. Mean intensities are quantified in arbitrary units (AU). Scale bars are 10 µm. Graph shows mean±SD. Significance (P-value) is from One-way ANOVA test and 2-tailed Student’s unpaired t-test. ALH means after larval hatching. Yellow arrowheads indicate NBs.

Since m3, m8, mβ, and mγ are the only members expressed in A3-A10 NBs prior to their death, we tested single deletion mutants of mγ (*Δmγ/Δmγ*), m3 (*Δm3/Δm3*), and a double deletion mutant of mβ and m3 (*Δmβm3/Δmβm3*)(which survived till late L3 stage) (70) (72), for the block of NB apoptosis. We observed that abdominal and terminal NBs apoptosis was blocked only in homozygous double deletion for mβ and m3 genes (*Δmβm3/ Δmβm3*) (Fig 3C-3D) at the LL3 stage. There were no surviving NBs in wild-type control, or in homozygotes for the single deletion mutants [for m3 (*Δm3/Δm3*) and mγ (*Δmγ/Δmγ*)] or the trans-heterozygotes (*Δmβm3/Δmγ, Δmβm3/Δm3, Δm3/ ΔE(spl)-C and Δmγ/ΔE(spl)-C*) (Fig 3E). Thereafter, we did individual and double RNAi knockdowns of E(spl)-HLH members from embryonic to LL3 stage (TS protocol: Fig. 3F’). We observed that while the single knockdown of m3, m8, mβ, and mγ did not show any surviving NBs, the double knockdown of different combinations of E(spl) HLH members (*mβ/mγ, m3/mγ, m8/mγ, m3/mβ*) consistently showed a block of NB apoptosis to various extents (Fig. 3F and Fig. S3E-S3H). A double knockdown of mβ with m5 (which do not express in NBs) did not show any block of NB apoptosis, thereby further corroborating the functional specificity of complex members in the CNS (Fig 3F and Fig. S3I-S3J). These results were consistent with previous reports showing that members of the E(spl)-HLH family function redundantly (73–75).

E(spl)-HLH members are TFs, and they can impact NB apoptosis either by transcriptionally regulating Hox, Grh, or members of the Notch signalling pathway, or by direct binding onto the apoptotic enhancer. Considering this, we checked the expression of Hox (Abd-B), Grh, and Cut in the double deletion for *E(spl)-mβ* and *m3* (*Δmβm3/Δmβm3*) and *Canton S* controls (at mid L3 stage) in A8-A10 NBs. Since Cut is a target of Notch signalling and E(spl)-complex (76) (61, 62), we expected that if mβ and m3 were regulating the members of the Notch signalling pathway in NBs, it would be reflected by a change in the Cut levels in the NBs. However, we did not observe any significant difference in the levels of Cut, Grh, and Abd-B in double deletion (*Δmβm3/Δmβm3*) (Fig. 3G-3I and Fig. S3K-S3Q) when compared to the controls. This suggested that these members most likely regulate NB apoptosis either by directly binding to the apoptotic enhancer or by regulating another (yet to be identified) factor involved in apoptosis. Considering that *E(spl)-mγ-GFP* expression and Notch signalling temporally increase before NBs apoptosis (56), we support the idea that E(spl)-C members probably work with Su(H) to temporally consolidate the impact of Hox/Grh and Notch signalling inputs on the apoptotic enhancer to execute cell death.

Collectively, the above results establish that members of the E(spl)-HLH (m3, mβ, m*γ* and m8) complex function redundantly in NBs, and are combinatorially required for executing NB apoptosis, and do not affect Hox, Grh and Notch pathway.

### Functionally conserved phosphatase activity of PP1α is required to promote NB apoptosis in vivo

The PP1α family of phosphatases is highly conserved across species, and *Drosophila* PP1-α87B is homologous to Human PPP1 (or hPPP1). Therefore, we wanted to assess functional conservation between the two orthologs and determine whether phosphatase activity was essential for promoting NB apoptosis in vivo. hPPP1 has a highly conserved Histidine residue at the 125^th^ position in its catalytic core, which is critical for its phosphatase activity (this residue corresponds to His-123 in *Drosophila* PP1α-87B) (Fig. 4A). Mutation of this Histidine to Glutamine is known to abrogate the phosphatase activity of the enzyme (63, 77–79). We tested this using an in vitro pNPP (p-NitroPhenylPhosphate) assay in which a catalytically active phosphatase cleaves pNPP to inorganic phosphate and pNP (estimated at 405 nm). We indeed observed that bacterially purified wild-type hPPP1 was enzymatically active while hPPP1^H125Q^ was not (Fig. 4B). Thereafter, we tested their capacity to dephosphorylate Su(H) specific synthetic phospho-peptide (^262^ALFNRLRS^P^QTVSTRY^276^; phosphorylated at S269) (9) using Malachite green (which absorbs at 655nm). Here, we observed that the wild-type hPPP1 could dephosphorylate the phosphopeptide at Ser 269, but the phosphatase-dead version could not (Fig. 4C). This established that hPPP1 could dephosphorylate Su(H) at Ser 269. This in vitro experiment was corroborated in vivo, wherein we observed that the block of NB apoptosis in A3-A10 segments caused by *PP1-α87B* knockdown (Fig. 4D and 4G) was rescued by temporally regulated overexpression (TS protocol: Fig. 4H) of hPPP1 (Fig. 4E and 4G), but not by hPPP1^H125Q^ (Fig. 4F-4G).

**Fig 4.**
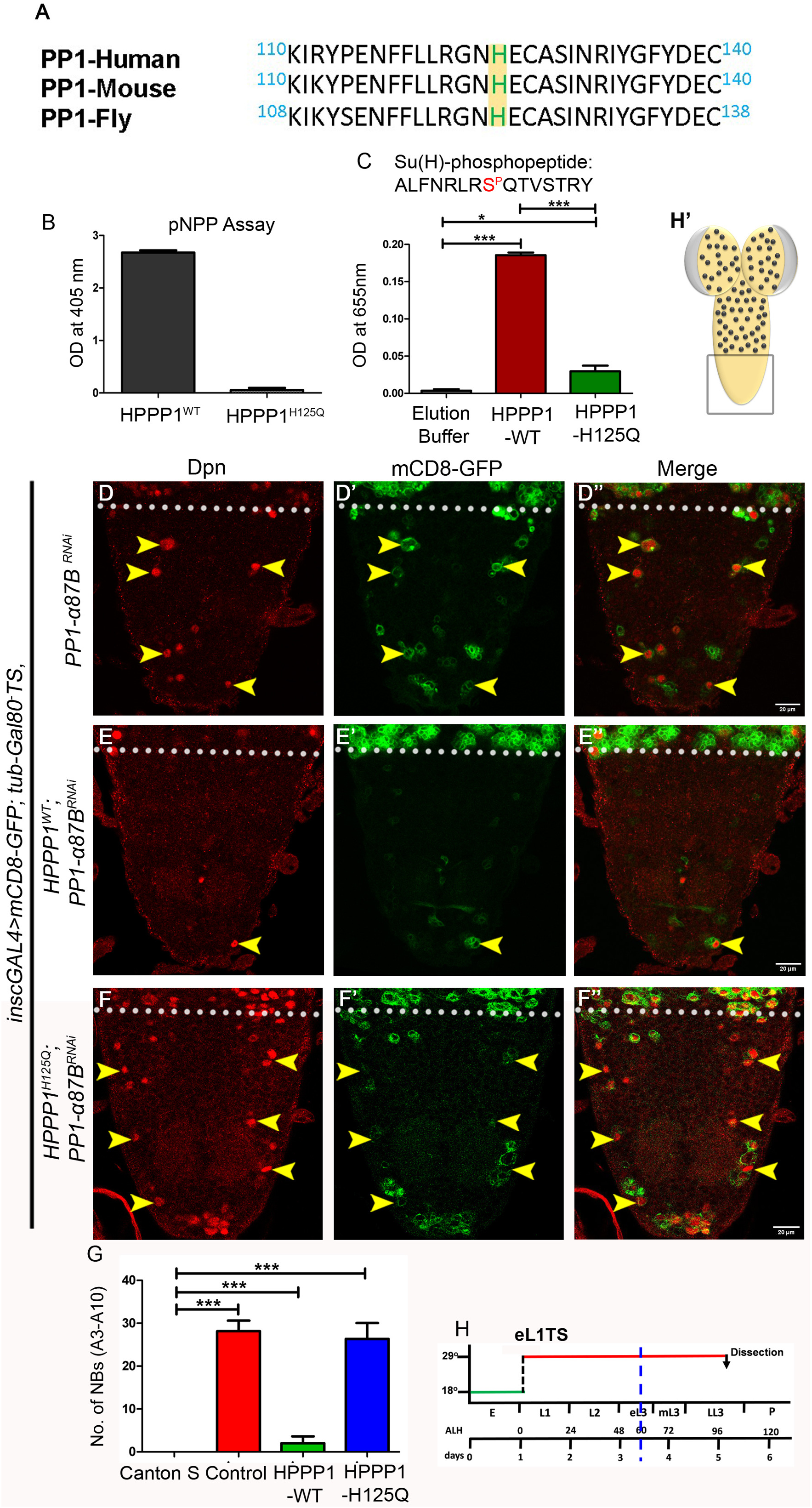
Functionally conserved phosphatase activity of PP1α is required to promote NB apoptosis and Notch signalling *in vivo*. (A) Alignment of a region from *Human, Mouse and Drosophila* PP1α, highlighting the conserved Histidine residue in green (H125 in *H. Sapiens* and *M. musculus;* H123 in *D. melanogaster*). (B) Graph showing that the H125Q mutation abolishes phosphatase activity of wild-type hPPP1 (n=3). The levels of p-NitroPhenol (pNP, estimated at 405nm) released by purified wild-type (hPPP1^WT^) and phosphatase dead (hPPP1^H125Q^) versions of human PPP1 by enzymatic cleavage of pNPP. The amount of pNP released from pNPP is indicative of the phosphatase activity of the enzyme. (C) Graph showing in vitro phosphatase assay establishing that hPPP1 can dephosphorylate Su(H) at Serine 269 (n=2). Graph quantifies the levels of released inorganic phosphate (detected by Malachite Green at 655nm) from the Su(H) specific phosphopeptide (*D. melanogaster:* ALFNRLRS^P^QTVSTRY) by bacterially purified hPPP1^WT^ and hPPP1^H125Q^. (D-F) Human ortholog of PP1-α (hPPP1) rescues *PP1-α87B* knockdown-mediated block of NB apoptosis. (D) *PP1-α87B* knockdown blocks NB apoptosis (*PP1-α87B-RNAi*: 28.14±2.47 NBs, n=7 VNCs, N=3). (E) The overexpression of wild-type human ortholog of PP1-α (hPPP1: 2.0±1.61 NBs, n=11 VNCs, N=3) rescues the block of NB apoptosis as evident by reduction in the number of Dpn-positive NBs. (F) The overexpression of phosphatase dead form of human PP1 (hPPP1^H125Q^: 26.33±3.72NBs, n=6 VNCs, N=2) is unable to rescue the block of NB apoptosis, suggesting the functional conservation of the role of PP1-α and its phosphatase activity in regulating Notch signalling. (G) Graph showing the number of surviving NB in A3-A10 segments for the *CantonS* controls and the genotypes shown in panels D-F. (H) Show the early L1 temperature shift (TS) protocols used in the experiment. (H’) Region of the CNS shown in all the panels of the figure is indicated by the box. All representative images shown are single confocal sections from LL3-stage female VNCs. Scale bars are 20 µm. Graph shows mean±SD. Significance (P-value) is from the One-way ANOVA test and 2-tailed Student’s unpaired t-test. ALH means after larval hatching. Yellow arrowheads indicate NBs. The white dotted line separates thoracic (T) and abdominal (A) segments of the VNC.

Together, these data suggest that PP1-α phosphatase activity is required to modulate Notch signalling by dephosphorylating Su(H) at Ser-269. The capacity of hPPP1A to dephosphorylate *Drosophila* Su(H) in vitro and to rescue *PP1α-87B* knockdown phenotype in vivo suggest that regulation of Notch signalling by PP1-α is likely to be conserved across species and may be widely used across the phyla during development.

### PNUTS functions as a regulatory subunit of PP1α in its modulation of the Notch signalling pathway

The catalytic subunit of PP1α functions as a holoenzyme and forms context-specific complexes with different regulatory subunits (PIPs). The PIPs are known to function as scaffolding proteins and can control in vivo functions of PP1-α by regulating its localisation (to a specific organelle), activity and target specificity (24, 80, 81). Therefore, one or more PIPs are expected to associate with PP1 in vivo to regulate its context-specific functions. We wanted to identify the PIP that worked with PP1-α to regulate the Notch signalling pathway in the context of NB apoptosis. To this end, we carried out an RNA interference screen in which we temporally induced the knocked down of different PIPs (TS protocol: Fig 5Q’) in larval NBs and scored for a block of NB apoptosis in A3-A10 segments of the larval CNS (Fig. 5Q). We tested 24 genes (**Table-1**) and observed that temporally controlled knockdown of *PNUTS* (**<u>P</u>**hosphatase 1 **<u>NU</u>**clear-**<u>T</u>**argeting **<u>S</u>**ubunit) by two independent RNAi lines (Fig. 5A-B: BDSC# 64538 and Fig. S5A and S5B, VDRC# 106862) resulted in block of NB apoptosis (Fig. 5B and Fig. S5B). Since these PNUTS RNAi lines have not been characterised previously, we tested them for the knockdown of PNUTS protein and RNA transcripts, and observed a significant reduction in the levels of PNUTS staining in thoracic NBs (Fig. S5G-S5I), and almost 80% reduction in mRNA levels (BDSC#64538; Fig S5F). The block of apoptosis observed in the case of *PNUTS* knockdown was also associated with a reduction in the levels of *E(spl)mγ-GFP* (Fig. 5C-5E) and Cut (Fig. S5C-S5E) in the surviving NBs in A3-A10 segments (compared to the *p35*-expressing control NBs), indicating a reduction in Notch activity in these cells, similar to what was seen in the case of *PP1-α87B* knockdown.

**Fig 5.**
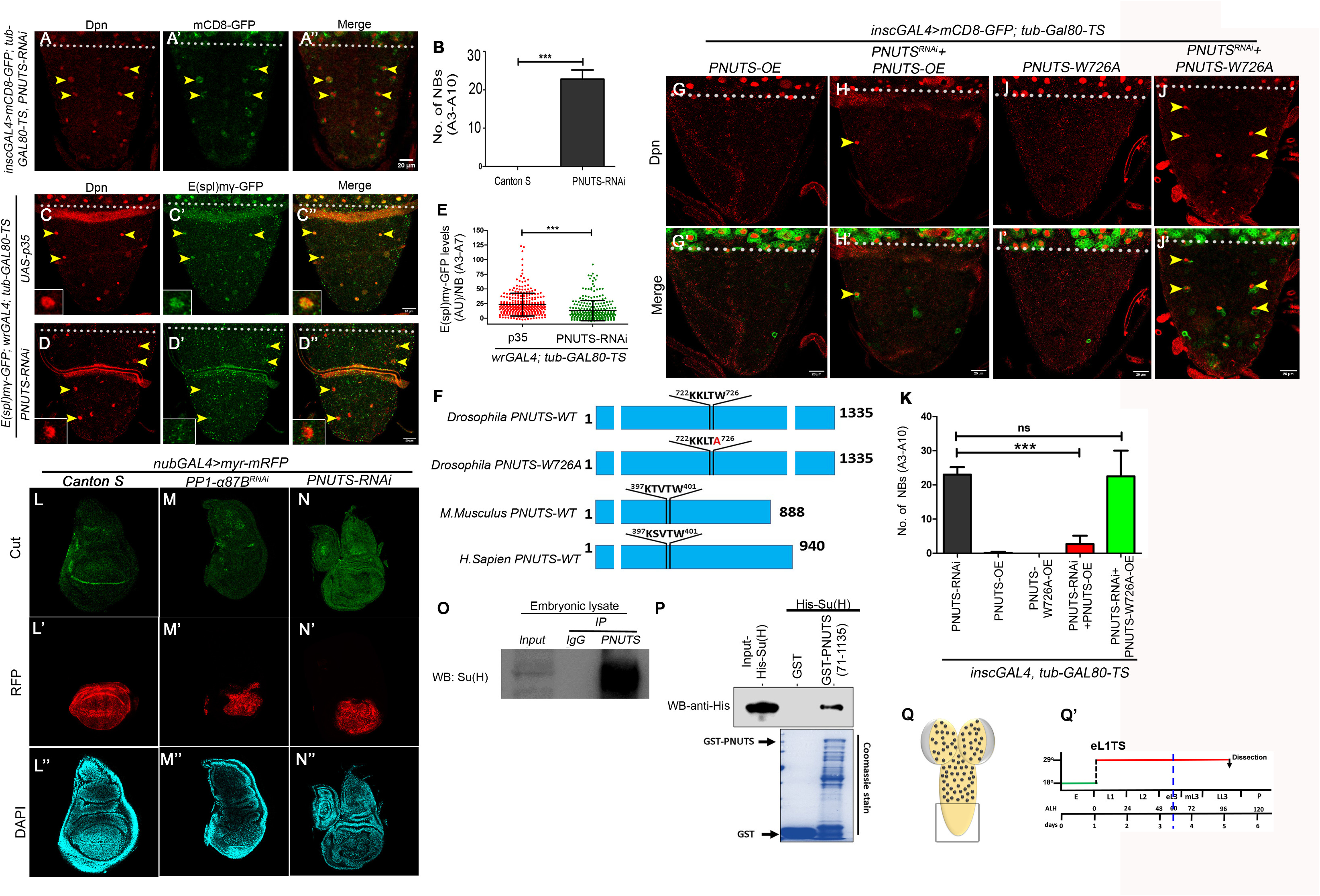
PNUTS functions as the regulatory subunit of PP1α to modulate the Notch signalling pathway. (A) Shows the surviving NBs (Dpn-positive cells) at the LL3 stage in A3-A10 segments of the larval VNC at the LL3 stage for *PNUTS* knockdown. (B) Graph comparing the number of surviving NB in A3-A10 segments for the *CantonS* controls (0±0 NBs, n=6 VNCs, N=3) vs *PNUTS* knockdown (23.00±2.19 NBs, n=6 VNCs, N=3). (C-D) Show that upon the knockdown of *PNUTS*, the levels of *E(spl)mγ-GFP* are reduced in the surviving NBs when compared to *p35* expressing controls blocked for NB apoptosis (TS: panel Q’). This suggests that *PNUTS* knockdown blocks NB apoptosis by affecting Notch signalling. (E) Graph comparing the intensities of *E(spl)mγ-GFP* in A3-A10 NBs in *p35* expressing control VNCs (23.21±19.39 AU, 300 NBs, n=12, N=3) versus VNCs with *PNUTS* knockdown (12.88±17.20 AU, 275 NBs, n=13, N=3). (F) Schematic of *Drosophila*, Mouse and Human PNUTS protein, highlighting the (conserved) putative PP1-binding motif (K/R-K-(S/T)-V/L-T-W). The schematic also shows a mutant version of *Drosophila* PNUTS with Tryptophan 726 replaced with an Alanine (PNUTS^W726A^), which abolishes its interaction with PP1α. (G-J) Shows the number of surviving NBs (Dpn-positive cells) in A3-A10 segments of the larval VNC of different genotypes at the LL3 stage. (G and I) Overexpression of PNUTS^WT^ (0.10±0.31 NBs, n=10Z, N=3) and PP1-α87B binding deficient version of PNUTS (PNUTS^W726A^) (0±0 NBs, n=8, N=3) does not block NB apoptosis. (H) Overexpression of wild-type PNUTS rescues the block of NB apoptosis resulting from PNUTS knockdown (2.66±2.44NBs, n=9 VNCs, N=3) as evident by a reduction in the number of Dpn-positive cells. (J) Overexpression of PP1-α87B-binding-deficient version of PNUTS (PNUTS ^W726A^) could not rescue the block of NB apoptosis (22.50±7.52 NBs, n=8 VNCs, N=3). These results establish the specificity of the PNUTS knockdown phenotype and show that PNUTS must interact with PP1-α87B to activate Notch signalling and execute NB apoptosis. (K) Graph showing the number of surviving NB in A3-A10 segments for the genotypes shown in panels A and G-J. (L-N) Shows that compared to controls, the Cut staining (Notch target) is compromised in the pouch of larval wing discs in the case of *PNUTS* (N) and *PP1-α87B* (M) knockdown. This suggests that PP1-α87B and PNUTS regulate Notch signalling in larval wing disc as well. Controls (*nubGal4>Canton.S*: n=8 wing disc, N=3); *PP1-α87B* (*nubGal4>PP1-α87B*-*RNAi*: n=15 wing disc, N=4); *PNUTS* (*nubGal4>PNUTS RNAi*: n=14 wing discs, N=3). (O) Western blot showing that antisera for PNUTS could pull down Su(H) from embryonic lysate while IgG could not, suggesting that PNUTS and Su(H) exist as a complex in vivo. (P) Bacterially purified GST-tagged PNUTS (71-1135) could pull down His-tagged Su(H) in vitro, implying that the interaction is direct. Coomassie Blue gel depicts the loading of the GST-tagged protein samples. Gels and blots shown are representative of 2 repeats. (Q) Region of the CNS shown in all the panels of the figure is indicated by the box. (Q’) Show the early L1 temperature shift (TS) protocols used in panels A, C-D, G-J. All representative images shown are single confocal sections from LL3-stage female VNCs. Mean intensities are quantified in arbitrary units (AU). Scale bars are 20 µm. Graph shows mean±SD. Significance (P-value) is from the One-way ANOVA test and 2-tailed Student’s unpaired t-test. ALH means after larval hatching. Yellow arrowheads indicate NBs. The white dotted line separates thoracic (T) and abdominal (A) segments of the VNC.

**Table 1:**
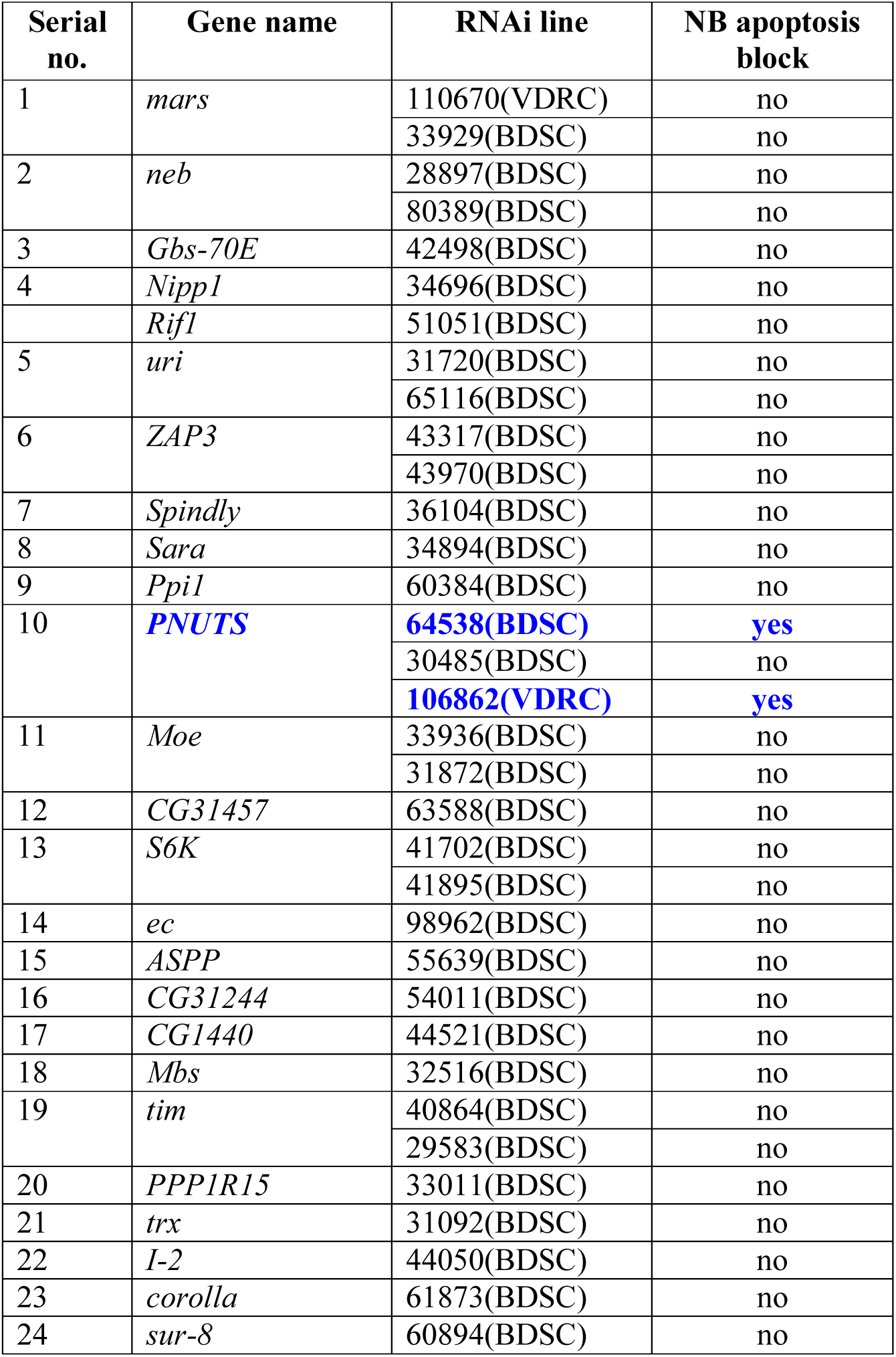
RNAi lines used for PIP screening.

Next, we wanted to confirm the role of PNUTS in NB apoptosis and check whether the regulation of Notch signalling by PNUTS is PP1-α87B dependent or independent. PNUTS has been proposed to function as a scaffold for PP1α and its substrate (26, 82) and contains a canonical PP1α-interacting motif K/R-K-(S/T)-V/L-T-W (26, 82–84) (Fig. 5F). In *Drosophila* PNUTS, the mutation of tryptophan to alanine at the 726^th^ position of the PP1α interaction motif (residues 722–726: KKLTW) (PNUTS^W726A^) abolishes its binding to PP1-α87B and reduces PP1-α87B function (26). Therefore, we performed an experiment to rescue the block of A3-A10 NB apoptosis (observed in PNUTS knockdown -Fig. 5A) by overexpressing (TS protocol: Fig. 5Q’) RNAi-resistant wild-type PNUTS (PNUTS^WT^) and a PP1-binding-deficient PNUTS (PNUTS^W726A^). We observed that while the PNUTS^WT^ rescued the block of NB apoptosis by reducing the number of Dpn-positive NBs in A3-A10 segments (Fig. 5H and bar-4 of Fig. 5K), the binding-deficient PNUTS (PNUTS^W726A^) failed to do the same (Fig. 5J and bar 5 Fig. 5K). Furthermore, only the overexpression of PNUTS^WT^ and PP1-binding-deficient PNUTS (PNUTS^W726A^) by themselves did not show a block of NB apoptosis (Fig. 5G, 5I and 5K). These experiments confirmed the role of PNUTS in the regulation of Notch signalling and established the specificity of the knockdown and PNUTS’s requirement to interact with PP1-α87B to regulate the Notch signalling pathway.

Since Notch signalling is important in multiple developmental contexts, we wanted to determine whether PP1α/PNUTS-mediated Notch regulation is specific to the CNS or extends to other cellular contexts as well. To address this, we examined the Notch-responsive Cut expression at the dorsal-ventral boundary of the wing disc (61) and observed that, like in NBs, Cut expression was dramatically repressed in the dorsal-ventral boundary of the wing disc in response to both *PP1-α87B* (Fig. 5M) and *PNUTS* knockdowns (Fig. 5N) compared to controls (Fig. 5L).

Lastly, we wanted to check whether PNUTS physically interacted with Su(H) in vivo. Here, we used an anti-PNUTS antibody and successfully pulled down Su(H) from whole-embryonic lysate (Fig. 5O). Thereafter, we performed an in vitro pulldown assay with bacterially purified His-tagged Su(H) and GST-tagged PNUTS and found that PNUTS pulled down Su(H), suggesting a direct physical interaction between the two proteins (Fig. 5P). This confirmed our notion that PP1-PNUTS functions as a complex to regulate Notch signalling by directly modulating Su(H) DNA binding activity.

Collectively, these results establish that PNUTS functions as a PIP for PP1α and helps it to modulate the Notch signalling pathway by dephosphorylating Su(H) at Serine 269, which is critical for executing NB apoptosis in the larval CNS.

### PP1-α regulates the Early to Late temporal transition of larval NBs

The Notch signalling pathway is known to regulate the Early to Late (E>L) temporal transition in approximately ∼ 20% of the Type I NBs in the central brain (59). This transition happens in the early third instar stage (60 hrs ALH, dashed blue line in Fig. 6Q) when NBs repress Imp and start expressing Syp (Fig. 1A’). Therefore, we wanted to test whether PP1-α/PNUTS regulation of the Notch pathway affected this transition. To this end, we focused on Type I NBs in the thoracic region (T1-A2 segments, Fig. 6Q’) of larval VNC and did an inducible knockdown for *Notch* from the eL1 stage to LL3 stage (TS protocol: Fig. 6Q). This experiment was done with two independent RNAi lines for Notch (BDSC 28981 and 33616). In both cases, we observed a block of E>L transition in the thoracic NBs with persistence of Imp (Fig. 6A’ vs 6B’ and bar 2 in Fig. 6J) and absence of Syp (Fig. S6A’ vs S6B’ and bar 4 in Fig. 6J) in approximately 15% of the NBs at the LL3 stage. This phenotype was similar but less penetrant than the phenotype reported in the central brain (59). Next, we knocked down *PP1-α87B* and observed persistence of Imp expression in 85% of thoracic NBs at the LL3 stage (Fig. 6A’ vs 6C’ and bar 2 in Fig. 6K), normally, all NBs should be Syp-positive by this time (bar 1 in Fig. 6L). This was further supported by the fact that only 50% of NBs had Syp expression in *PP1-α87B* knockdown (bar 2 in Fig. 6L and Fig. S6C’). This suggested a role for PP1-α in executing this transition in the majority of NBs. These *PP1-α87B* knockdown associated E>L transition defect phenotypes were completely rescued by the overexpression (TS protocol: Fig. 6Q) of an RNAi-resistant version of *PP1-α87B* (Fig. 6D’ and Fig. S6D’) or the human ortholog of this PP1-α (hPPP1) (Fig. 6E’ and Fig. S6E’), both of which resulted in ∼100% of NBs expressing Syp (bar 3 and 4 Fig. 6L) and very few NBs (bar 3 and 4 Fig. 6K) expressing Imp at the LL3 stage. However, the catalytically dead version of the human ortholog (hPPP1^H125Q^) was unable to rescue the phenotype (Fig. 6F’ and Fig. S6F’), with ∼85% NBs still expressing Imp (bar-5 in Fig. 6K) and only ∼45% NBs expressing Syp (bar-5 in Fig. 6L) at the LL3 stage. The rescue experiment with PP1-α87B and hPPP1 establishes the specificity and the possible functional conservation of PP1α in executing the E>L transition. The absence of the rescue of the E>L transition in hPPP1^H125Q^ establishes the importance of the phosphatase activity of PP1α in mediating this effect.

**Fig 6.**
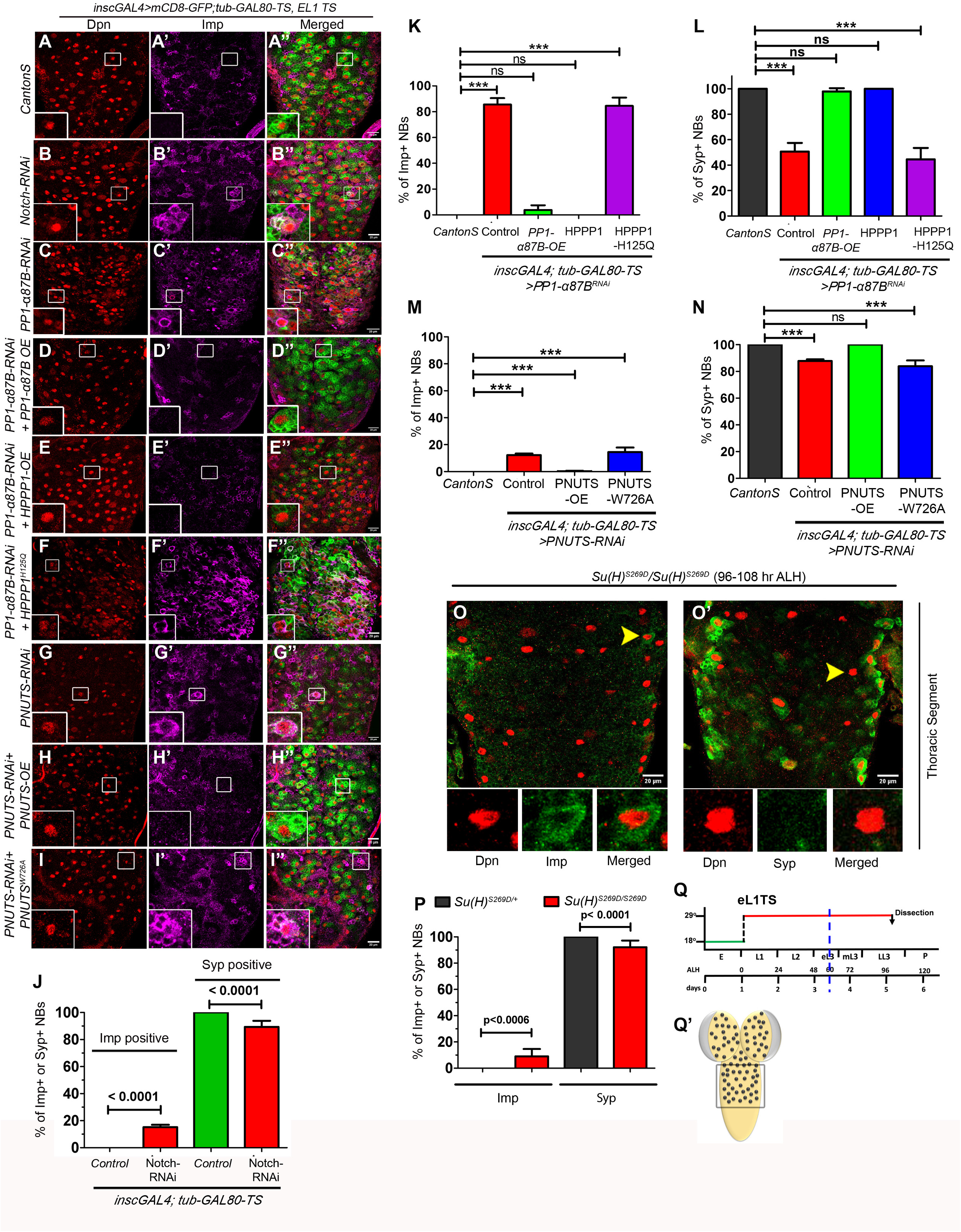
PP1-α87B/PNUTS controls the Early-to-Late competence switching of the thoracic NBs, partly through Notch pathway regulation. (A-I) Shows the thoracic segments of LL3 stage larvae of the indicated genotypes. Brains are immunostained with antisera for early temporal factor Imp (Magenta) and NB marker Dpn (Red). (A) The NBs in the control (*CantonS*) VNCs do not express Imp by the LL3 stage (0% NBs, n=9 VNCs, N=3), but upon *Notch* knockdown (B), 15% of thoracic NBs (15.18±1.78% NBs, n=9, N=3) retain Imp expression, indicating their inability to undergo the competence switch. (C) Knockdown of *PP1-α87B* results in the majority of NBs retaining Imp expression (85.63±4.95 % NBs; n=8 VNCs; N=3) at the LL3 stage. This defect is rescued by overexpression of *Drosophila PP1-α87B* (3.75±3.69% NBs; n=7 VNCs; N=3) (D), and its human ortholog (hPPP1: 0% NBs, n=7 VNCs, N=3) (E), but not by phosphatase dead version of the same (hPPP1^H125Q^: 84.50± 6.433% NBs, n=10, N=4), establishing the specificity of the phenotype and suggesting a cross phylum conservation of the role of PP1-α in regulating competence switching of the NBs. (G) Shows that upon knockdown of *PNUTS*, 12% of thoracic NBs (12.18 ±1.19 % NBs, n=9 VNCs, N=3) retain Imp expression, indicating their inability to undergo the competence switching. (H-I) This defect is rescued by overexpression of wild-type *Drosophila PNUTS* (*PNUTS-OE*: 0.26±0.37% NBs, n=8 VNCs, N=3) (H), but not by the PP1-α-binding-deficient form of *PNUTS* (*PNUTS^W726A^*:14.50±3.36% NBs, n=9 VNCs, N=3), establishing the specificity of the phenotype and suggesting that PNUTS need to interact with PP1α to execute the competence switching. (J) Graph showing the percentage of thoracic NBs expressing early factor Imp and late factor Syp (Fig. S6A-S6B) for *CantonS* control and *Notch* knockdowns shown in panels A and B. (K-N) Graphs showing the percentage of thoracic NBs expressing early factor Imp (K and M) and late factor Syp (Fig. S6C-S6I) (L and N) in the case of the genotypes shown in panels C-I. (O-P) Shows the thoracic segment of the larval VNC homozygous for *attP-Su(H)* allele, where Serine 269 is replaced with an Aspartate (*attP-Su(H)^S269D^/attP-Su(H)^S269D^*) (108 hrs ALH). Approximately 9% of NBs from these VNCs retain Imp staining in LL3 stages (Imp-positive NBs: 9.03±5.59% NBs, n=10 VNCs, N=4), congruent to that approximately 8% of the NBs were unable to express Syp, and the remaining 92% were positive for Syp (Syp-positive NB: 92.09±5.15, n=8 VNCs, N=4). (P) Graph showing the percentage of thoracic NBs expressing early factor Imp and late factor Syp in the case of heterozygous control VNCs (*attP-Su(H)^S269D^/+*) and homozygous test VNCs (*attP-Su(H)^S269D^/attP-Su(H)^S269D^*) shown in panels O and O’. This suggests that PP1α regulates competence switching in 85% of the NBs; in 15% of the cells, PP1α functions by regulating Notch signalling. Of which in 9% of cells PP1α work by dephosphorylating Su(H) at Serine 269, in the remaining 6% of the NBs, PP1α modulates Notch signalling by dephosphorylating an unidentified residue to regulate competence switching. (Q) Show the early L1 temperature shift (TS) protocols used in panels A-I. (Q) Region of the CNS shown in all the panels of the figure is indicated by a box. The insets in panels A-I are the magnified view of the NBs indicated by white boxes. All representative images shown are single confocal sections from LL3-stage female VNCs. Scale bars are 20 µm. Graph shows mean±SD. Significance (P-value) is from the One-way ANOVA test. ALH means after larval hatching.

Thereafter, we knocked down *PNUTS* (TS protocol: Fig. 6Q) to test whether it was the regulatory subunit required by PP1-α for executing the E>L transition in thoracic NBs. Here, as with Notch knockdown, *PNUTS* knockdown also resulted in approximately 12% of NBs showing a block of the E>L transition (Fig. 6A’ vs 6G’) and were Imp-positive (bar 2 in Fig. 6M), compared to 85% cells that showed a block of the transition in the case of PP1-α knockdown at the LL3 stage. In support of this, 13% of the NBs did not express Syp at the LL3 stage (bar 2 in Fig. 6N; Fig. S6G’ vs S6A’). By this time, all the NBs are expected to be Syp-positive (bar 1, Fig. 6N). This was further supported by the fact that *PP1-α87B* and *PNUTS* knockdowns (TS protocol: Fig. 6Q) resulted in a reduction in the levels of *E(spl)mγ-GFP* in the Type I NBs of the thoracic segment (Fig S6J-S6M) in the LL3 stage. These observations supported the idea that NBs, which rely on the Notch pathway for the E>L transition, may use PNUTS as a regulatory subunit for PP1-α. Next, we wanted to assess the specificity of the E>L transition defect observed in the case of *PNUTS* knockdown and whether the ability to interact with PP1α was critical for the transition. To this end, we attempted to rescue the transition defect using an RNAi-resistant version of *Drosophila PNUTS* and a PP1-α-binding-deficient PNUTS. Here we observed that the overexpression of RNAi-resistant version of *Drosophila PNUTS* completely rescued the knockdown phenotypes (TS protocol: Fig. 6Q), wherein 0% NBs were Imp-positive (Fig. 6H’ and bar-3 of Fig. 6M) and all 100% NBs had transitioned into Imp-negative/Syp-positive state at the LL3 stage (Fig. S6H’ and bar-3 of Fig. 6N). However, the overexpression of the PP1-α binding-deficient version of PNUTS (PNUTS^W726A^) could not rescue the E>L transition defect (Fig. 6I’) with ∼18% of NBs being Imp-positive (bar 4 of Fig.6M) (as against 0% NBs in wild-type controls) at LL3 stage, and only ∼85% of the NBs being Syp-positive (bar-4 of Fig 6N and Fig. S6I’) (as against 100% NBs in wild type controls) similar to what was observed for PNUTS knockdown. This suggested that PNUTS’s capacity to rescue the E>L transition defect critically relies on its ability to interact with PP1-α.

In the context of NB apoptosis, we had observed that PP1-α mediated its effect on Notch signalling by dephosphorylating Su(H) at Serine 269. We tested whether the same residue was relevant to the E>L transition. This was done using an attP replacement allele of Su(H) (*attP-Su(H)^S269D^*) (64). We assessed the control heterozygous larvae of the attP allele (*attP-Su(H)^S269D^*/+) at the LL3 stage (96 hrs ALH). We observed that they did not show any E>L transition defect, with all the 100% NB transitioning into Syp-positive state, with no Imp-positive NBs remaining (bars 1 and 3 in Fig. 6P). The age-matched homozygous larvae of the attP allele (*attP-Su(H)^S269D^*/*attP-Su(H)^S269D^*) at 96 hrs ALH showed a significant developmental delay and were found to be in the L3 stage (had L3 spiracles, which had not everted out from the body wall) and were dead 16 hrs later. In this case, we analysed the VNCs at 108 hrs ALH and observed a block of E>L transition in 9% of the NBs (Fig. 6O and bar-2 in Fig. 6P), which were Imp-positive (as opposed to 0% imp-positive NBs found in controls) and had failed to activate Syp. Similarly, 90% of the NBs were Syp-positive (as opposed to 100% in controls) (Fig. 6O’ and bar-4 in Fig 6P).

These results collectively suggest that PP1-α/PNUTS-mediated dephosphorylation of Su(H) at Ser 269 is important in 9% of the NBs to execute the E>L transition. The remaining 6% of the NBs need to be dephosphorylated at other (yet to be identified) residue/s to execute the E>L transition.

Since E(spl)-HLH complex members m3 and mβ were primarily responsible for executing NB apoptosis downstream of the Notch signalling pathway, we wanted to check if the same two members were responsible for executing E>L transition as well. For this, we analysed the homozygotes of the double deletion for m3 and mβ (*Δmβm3/Δmβm3*) and found only a very weak effect (Fig. S6N). At the LL3 stage, none of the NBs showed Imp expression, and almost 95% of the NBs were found to be Syp-positive (Fig. S6N). This suggests that, unlike NB apoptosis, E(spl) m3 and mβ do not play a role in E>L transition in NBs.

Collectively, our results suggest that PP1-α mediates E>L transition in 85% of the Type I NBs in larval VNC (Fig. 7A). In 15% of these cells, this transition is mediated by PP1-α/PNUTS-mediated regulation of the Notch signalling pathway (Fig. 7A), which is partially executed (in 9% of the NBs) through dephosphorylation of Su(H) at Ser-269 (Fig. 7B). We also find that the Notch pathway does not rely on E(spl) m3 and mβ members to execute the E>L transition in NBs. The result also suggested that the PP1-α-mediated E>L transition in the majority of NBs uses a Notch-PNUTS independent mechanism to execute it.

**Fig. 7.**
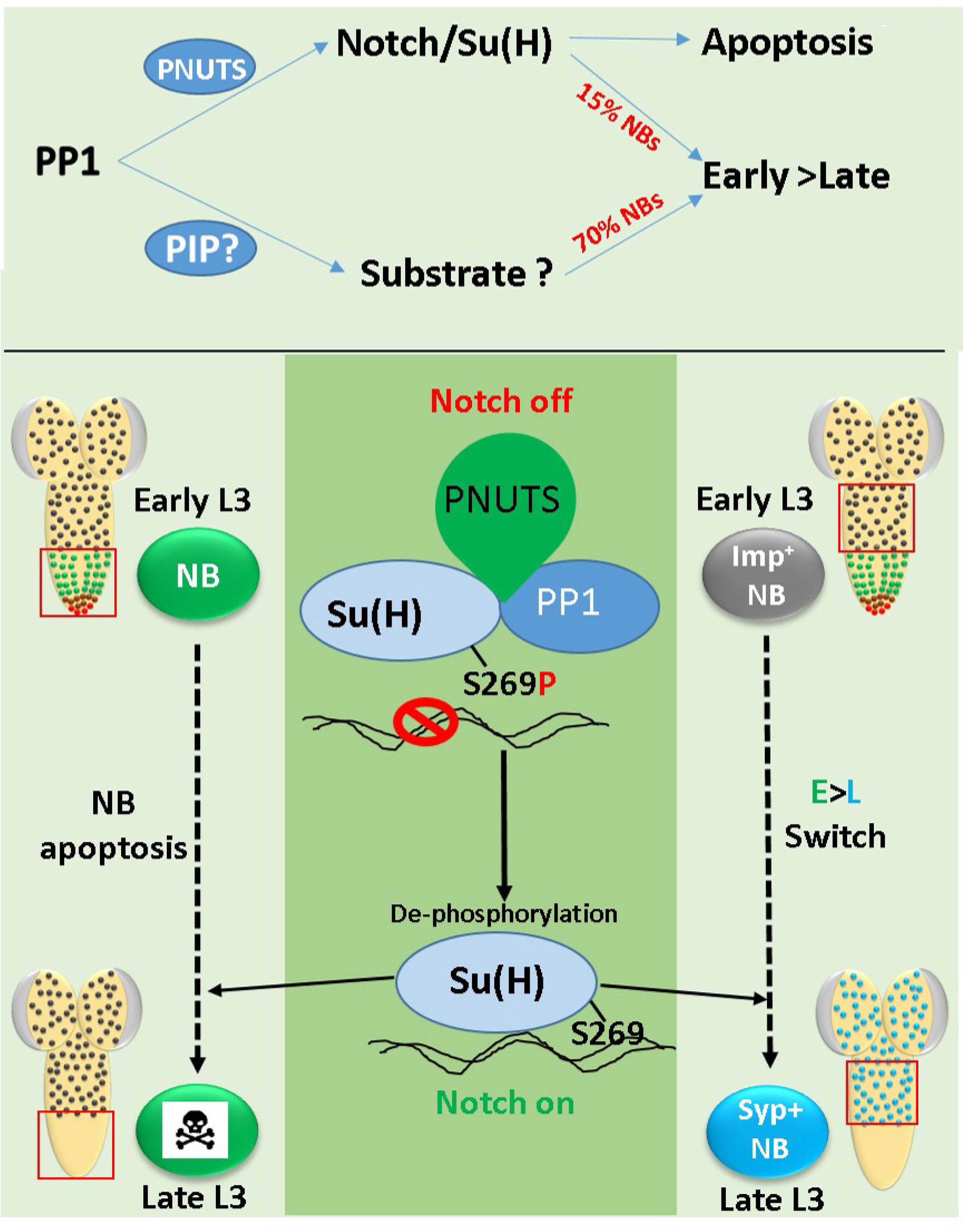
Graphical Summary. (A) PP1-α and its regulatory subunit PNUTS are required for apoptosis of all A3-A10 NBs in the VNC. PP1-α is also required for competence switching (E>L transition) in 85% of the thoracic NBs; a subset of ∼15% of these cells use Notch signalling to execute the competence switch, and in these cells, PP1-α uses PNUTS as its regulatory subunit. In the remaining ∼70% of the NBs, the mechanism of PP1-α-mediated competence switching and the regulatory subunit used by PP1-α are unknown. (B) Mechanistically, PP1-α/PNUTS dephosphorylate Su(H) at Serine 269, thereby restoring its DNA binding activity and activating downstream targets of Notch to execute apoptosis of NBs in A3-A10 segments of the CNS. This Serine 269 dephosphorylation is also repurposed as a regulator of the competence switching in ∼9% NBs (which use the Notch signalling pathway for E>L transition).

## Discussion

Our study identifies PNUTS as the regulatory subunit of PP1-α87B in the context of brain development and neurogenesis, and provides the first evidence of the two playing a crucial role in neurogenesis by modulating the activity of one of the most fundamentally important neurogenic pathways (Notch signalling). We show that PP1-α87B/PNUTS positively regulate Notch signalling at the level of posttranslational modification of Su(H) to facilitate developmental neurogenesis by executing two distinct physiological events in the NBs located in different regions of the larval CNS (Fig. 7B). In the first case, PP1-α/PNUTS dephosphorylate Su(H) at a conserved Serine residue (Serine-269) to regulate apoptosis of the NBs, thereby regulating the morphogenesis in the abdominal and terminal segments of the larval VNC. In the second case, we find that this posttranslational modification by PP1-α/PNUTS is also repurposed to partially regulate the early-to-late competence switch in a subset of thoracic NBs that use the Notch pathway to execute the switch (Fig. 7B). More specifically, we find that PP1-α is required to execute the early-to-late competence switch in the majority (∼85%) of NBs in the thoracic region of the VNC. A subset of these NBs (∼15% of the total 85%) use the Notch signalling pathway to execute the competence switch. In these 15% cells, PP1-α/PNUTS regulates Notch signalling by dephosphorylating Su(H) at Serine 269 in ∼9% of the NBs to execute competence switching. In the remaining 6% of cells, PP1-α/PNUTS dephosphorylate another unidentified residue on Su(H). This raises two important questions: the first is which residue(s) on Su(H) (other than Serine269) is/are responsible for executing the PP1-α/PNUTS-mediated competence switching in the remaining 6% of the NBs. The second question is how PP1-α regulates the competence switching in the remaining majority of NBs, and which regulatory subunit(s) facilitate this function, and what are the targets of PP1-α in these cells. We also found that the regulation of the Notch pathway by PP1-α/PNUTS was extendable to a different context of the wing disc tissue as well. Considering that this modulation of the Notch pathway critically relied on PP1-α phosphatase activity and that human PPP1 could rescue the PP1-α knockdown phenotypes, we believe this regulation is likely conserved across species during development.

Notch signalling is known to play an important role in neurogenesis (1, 59, 85, 86) and, therefore, is expected to be regulated at multiple levels (1, 5, 6). Upon signalling activation, NICD is cleaved, translocates into the nucleus, and binds to Su(H), altering its conformation from a repressor complex to a transcriptionally active complex. Therefore, the DNA-binding ability of Su(H) is crucial for Notch signal transduction. A previous study showed that mutation of a highly conserved Serine 269 to Aspartate (S269D) dramatically reduced its DNA binding (8). Here we show that PP1-α/PNUTS binds Su(H) and dephosphorylates it at Serine 269, thereby restoring its DNA-binding activity and contributing to signal transduction. Expectedly, the overexpression of the phospho-mimetic version of Su(H) (UAS-Su(H)^S269D^) or the attP replacement allele of Su(H) with substitution of Serine 269 by Aspartate (*attP-Su(H)^S269D^*) reduces Notch signalling in the abdominal and terminal NBs, thereby blocking the NB apoptosis. This is further supported by the fact that knockdown of *PP1-α87B* was rescued only by the overexpression of the phospho-dead version of Su(H) (Su(H)^S269Q^) but not by the phospho-mimetic version (Su(H)^S269D^). This supports the following two observations: first, that PP1-α87B acts upstream of the Su(H) activation complex; second, the phosphorylation event locks Su(H) into a repressive conformation (inhibiting its DNA binding and blocking the activation of downstream targets). This repressive conformation or Notch-Off condition is relieved either by dephosphorylation of phospho-Serine 269 or is perceived to be relieved by Su(H)^S269Q^ (which is a phospho-dead version and possibly mimics the dephosphorylated Su(H)) and therefore can rescue the phenotype, probably by forming an activated complex (with coactivator Mastermind). This raises an important question about the utility of this additional layer of regulation in the Notch signalling pathway. The Notch pathway has multiple ligands that elicit distinct signalling dynamics, leading to different downstream transcriptional programs and, therefore, altering the strength of signal transduction and resulting in distinct cell fates (87, 88). For example, Dll1 and Dll4 are known to transduce pulsatile versus sustained signalling downstream of the Notch receptor. Considering our results and previous reports (8, 9, 17), we believe that phosphorylation of Su(H) to abrogate Notch signal transduction and PP1-α-mediated dephosphorylation to reactivate it could be a molecular strategy to rapidly “switch on” and “switch off” the pathway in the responding cell. There are previous reports documenting the need for a rapid downregulation of Notch signalling in the vertebrate immune and nervous system development (89, 90). More specifically, in *Drosophila*, existing work has shown that during a parasitic wasp infestation (an immune challenge), Notch signalling needs to be downregulated to alter the cell repertoire required for wasp egg encapsulation. This rapid downregulation of the Notch pathway is carried out by Pkc53E-mediated phosphorylation of Su(H) at Serine 269, which reduces Su(H) DNA-binding activity (9). Understandably, PP1-mediated dephosphorylation would be expected to restore the Notch activity once the immune challenge has been neutralised, allowing normal hematopoiesis to resume. However, the molecular trigger and the kinase required to phosphorylate Su(H) in the context of neurogenesis are yet to be identified. We also showed that HLH members of the Notch-responsive E(spl) complex are crucial for promoting NB apoptosis. However, we did not find the same members of E(spl) complex to be critical for competence switching, suggesting a context-specific deployment of E(spl) complex members downstream of the Notch pathway in NBs to execute distinct physiological processes during neurogenesis. To summarise, our study identifies an important, unexplored molecular link between the Notch signalling pathway and a ubiquitously expressed and physiologically important enzyme, Protein Phosphatase 1, and elucidates how they together regulate the spatiotemporal development of the CNS.

## Materials and Methods

### Fly stocks and husbandry

The following fly lines were used: *Canton S* (BDSC-6349), *PP1-α87B RNAi* (BDSC-32414, v35024), *PP1-α87B^87Bg-3e1^ PP1α-96A^2^*/TM6B, Tb^1^(BDSC-23966), *attP-Su(H)^S269D^*/SM6CyO-GFP (gifted by A. Nagel) (64), *UAS-p35* (DGRC-108019), *UAS-PP1-α87B-OE* (RNAi resistant line-this study), *worniu-GAL4* (gifted by C. Doe), *F3B3-lacZ* (54), *E(spl)mγ-GFP* (60), *UAS-dcr2; inscGAL4 UASmCD8-GFP and UAS-dcr2; inscGAL4 UASmCD8-GFP; tub-GAL80^ts^* (gifted by J. A. Knoblich, (91)), *nub-GAL4.K, UAS-myr-mRFP* (BDSC-63148). *UAS-HA-Su(H)^WT^* (this study), *UAS-HA-Su(H)^S269D^* (this study) *UAS-HA-Su(H)^S269Q^* (this study), *UAS-HA-Su(H)^R266H^* (this study), *FRT-82B-E(spl)32.2* (BDSC-52011), *ΔE(spl)-m3^CR1^* (70), *ΔE(spl)-mγ* (70, 72), *ΔE(spl)-m3^CR1^mβ^CR1^* (*E(spl)*-*complex* member CRISPR deletions gifted by Schweisguth) (70), *E(spl)-m3-RNAi* (BDSC-25977), *E(spl)-mβ RNAi* (BDSC-26202), *E(spl)-m8 RNAi* (BDSC-26322), *E(spl)-mγ RNAi* (BDSC-25978), *E(spl)-m5 RNAi* (BDSC-51466, 26201), *UAS-hPPP1^WT^* (BDSC-64394), *UAS-hPPP1^H125Q^* (this study), *PNUTS RNAi* (BDSC-64538, VDRC-106862), *Notch RNAi* (BDSC-28981), *UAS-PNUTS-OE* (RNAi resistant line-this study), *UAS-PNUTS^W726A^-OE* (RNAi resistant line-this study), *elav[C155]-GAL4*, *UAS-mCD8-GFP*, *hsflp1*, *w* (BDSC,-5146), *yw*; FRT82B *tub-GAL80-LL3* (BDSC-5135).

All the fly stocks and crosses were maintained at 25°C unless mentioned otherwise. For experimental fly crosses, a 4-6 hrs window was kept to collect eggs, and the eggs were thereafter reared at 18° C or 29°C, depending on the experimental requirements. Larval staging was done based on number of hours after larval hatching (ALH) and appropriate stages were identified by stage specific larval spiracles. For early L1 stage specific knockdown or overexpression (OE) experiments, after egg collection, larvae were kept at 18°C for 48-56 hours and thereafter shifted to 29°C (early L1 temp shift-eL1TS) and reared for 92-96 hours (Late L3 stage). For experiments with *E(spl)-complex* single and combination mutants (Δm3, Δmγ, Δm3mβ), after 6-8 hours of egg collection, larvae are reared at 25°C till the late L3 stage.

### Clonal analysis

MARCM clones were generated as per previously described protocol (92). Briefly, after larval hatching, in early L1 stage (8 hrs ALH) larvae were heat shocked at 37°C for 1 hr. Thereafter, additional single or two heat shocks were given in 12 hour intervals. Larvae were further aged till late L3 stage at 25 °C, and dissections were performed for immunohistochemistry at approximately 96 hrs ALH.

### Immunohistochemistry and image acquisition

Female Late L3 larval brains were dissected in PBS and fixed in 4% paraformaldehyde in PBST (0.1% Triton-X in 1X PBS) for 30 minute in room temperature. After washing three times for 10 min each with PBST, brains were incubated with primary antibodies and kept overnight at 4°C. Next day, after three washes with PBST, brains were incubated with secondary Abs with appropriate dilution in PBST for 2 hours in room temp. After secondary Ab staining, brains were again washed with PBST for three times for 10 min each. Depending on the experimental needs, the following primary Abs were used: rabbit anti-Dpn (1:2000) (54); mouse anti-Grh (1:2000), mouse anti-Abd-A (1:1000), mouse Abd-B (1:500, 1A2E9, DSHB), mouse anti-NICD (1:50, C17.9C6, DHSB), mouse anti-Imp (1:1000, this study), mouse anti-Syncrip (1:1000, this study), mouse anti-Cut (1:20, 2B10, DHSB), chicken anti-GFP (1:2000, ab13970, Abcam), chicken anti-β-gal (1:2000, ab9361, Abcam), chicken anti-HA (1:500, ab9111, Abcam), mouse anti-Su(H) (1:500, SC-398453, Santa-Cruz), mouse anti-PNUTS (1:500, this study). Secondary antibodies conjugated to Alexa fluorophores from Molecular Probes were used: Alexa Fluor 405 (1:250); Alexa Fluor 488 (1:500); Alexa Fluor 555 (1:1000); and Alexa Fluor647 (1:500). Thereafter, brains were washed with PBST and mounted with ventral side up in 70% glycerol and images were acquired using ZEISS LSM 700 and 900 inverted confocal microscope.

The details of the PP1-α87B, PNUTS, Imp and Syncrip antibody generation are given in the supplementary material and methods.

### Image analysis

All images were processed and analysed using ImageJ/fiji. Scale bars are shown in the respective figure legends. All representative images shown are single confocal sections from late L3 stage female VNCs except for Fig. S3 (which shows both mid and late L3 stage VNCs). The graphs show mean±SD. Total number of surviving NBs were counted manually using the multipoint toll. To measure the mean fluorescence intensity, a circle around the Dpn positive NB were drawn at best focus using the “Oval selection tool” in ImageJ and then measured the intensity of respective channels. The calculated intensities were subtracted by background mean intensity from neuropil or the nearby region of the respective channel. Mean intensities are quantified in arbitrary units (AU). Percentage of Imp and Syncrip expressing thoracic NBs in *PNUTS* RNAi and UAS-Su(H)^S269D^ overexpression were calculated by manually counting the total thoracic NBs followed by Imp or Syp positive NBs. In case of *PP1-α87B* knockdown and related rescue experiments, across the z stack 20 random Dpn+ NBs were marked using the multi-point tool and further checked for the Imp/Syp staining to calculate the percentage of Imp/Syp expressing NBs. Microsoft Excel and GraphPad Prism were used for the data analysis. Significance (P-value) is from One-way ANOVA test and 2-tailed Student’s unpaired t-test. In ANOVA, Tukey’s post hoc test was used to obtain the P values. Unless otherwise mentioned at least three technical replicates were done for all the experiments.

### *In vivo* pulldown assays with whole embryo

Canton-S embryos were collected for 3-days and then aged till stage 15-16. After harvest, embryos were dechorionated and lysed using a Dounce homogenizer in lysis buffer containing 150 mM NaCl, 50mM HEPES (Ph 7.5),1 mM EDTA, 2.5mM MgCl2 0.2% Triton-X, 1mM PMSF and protease Inhibitor cocktail. The resultant protein extract was subjected to centrifugation and collected supernatant was further subjected to preclearing by incubating with Protein-A sepharose beads for 2 hrs. Resultant precleared lysate was then incubated overnight at 4°C, with control IgG and appropriate primary antibodies bound to Protein-A bead. Mouse anti-PP1 (this study), mouse anti-PNUTS (this study) and mouse anti-IgG (Jackson Laboratories) was used for IP. 50 μg of protein was used as input. After washing, bead-bound proteins were separated by 10% denaturing SDS-PAGE and transferred onto PVDF membrane. After blocking, membrane was incubated with rat anti-Su(H) Ab (1:2000, MABE982, Merck) overnight at 4°C. Further, HRP conjugated secondary antibody Peroxidase AffiniPure Goat Anti-Rat IgG (H+L) (1:10,000, 112-035-003; Jackson Immunological Research Laboratory, USA) were used. Visualization was carried out by femtoLUCENT™ PLUS-HRP Chemiluminescent detection kit (G Biosciences #786003) in Intelligent Image Quant LAS 500.

### *In vitro* GST-pulldown assay

The following proteins were used for *in vitro* pulldown: GST only, GST-PNUTS (71–1135), 6X-His-Su(H). GST-PNUTS (71–1135) was used for testing PNUTS and Su(H) interaction, owing to the poor expression often seen in the case of full-length PNUTS. In brief, bacterial cultures expressing full length/truncated GST-tagged and His-tagged proteins were induced for 5 h with 0.5 mM IPTG at 18°C. After sonication, lysates were treated with Micrococcal Nuclease (MNase) at 30°C for 10 mi (Nguyen and Goodrich 2009). Bead-bound GST proteins were incubated separately with an equal amount of His-tagged protein lysates for 2 hr at 4°C. After three washes bead-bound proteins were separated by denaturing SDS-PAGE and then transferred onto PVDF membranes. The membrane was further blocked in 5% milk in Tris-buffered saline with 0.1% Tween 20 (TBST). Primary antibody-mouse anti-His (H1029, Sigma-Aldrich) was diluted 1 in 5,000 in TBST, and the blot was incubated overnight at 4°C. The following HRP conjugated secondary antibody (Peroxidase-AffiniPure Rabbit Anti-Mouse IgG + IgM (H + L) (1:10,000, 315-035-048; Jackson Immunological Research Laboratory, USA) was used. Visualization was carried out by enhanced chemiluminescence detection. Representative blot from at least 3 repetitions of the experiments are shown in the figures.

### pNPP Phosphatase assay

Phosphatase enzymatic activity of *Drosophila* PP1-*α*87B and human wildtype hPPP1 and enzymatic dead hPPP1^H125Q^ was measured using the universal phosphatase substrate pNPP (MedChemExpress). In brief, bacterially purified 6X-His tagged PP1-*α*87B, hPPP1-WT and hPPP1^H125Q^ were incubated with 2.5 mM pNPP in phosphatase reaction buffer (10 mM HEPES, 150 mM NaCl, 1 mM MnCl2) at 37°C for 45 min. The released pNitroPhenyl (a byproduct of pNPP cleavage by the Phosphatase) was measured using a spectrophotometer at 405 nm.

### *In vitro* Phosphopeptide Phosphatase assays

Briefly, bacterially purified wild type PP1-α87B, hPPP1 and enzymatic dead hPPP1^H125Q^ were incubated with 100 µM phosphoserine peptide (generated by Biomatik) with the sequence ALFNRLR(pS)QTVSTRY, corresponding to the S296 region of Su(H) in phosphatase reaction buffer (10 mM HEPES, 150 mM NaCl, 1 mM MnCl2) at 37°C for 1hr. The amount of released inorganic phosphate was measured calorimetrically using the malachite green reagent kit (Malachite Green Phosphate Assay Kit, CST). The Malachite green assay was performed according to the manufacturer’s protocol where 25µl reaction mixture from the phosphatase reaction was transferred to a microtiter plate and 100µl of malachite green reagent was added to each well. After 20 minutes of incubation at room temperature, absorbance was measured at 655 nm using a microplate spectrophotometer.

## Supplementary Figure Legends

**Fig S1:**
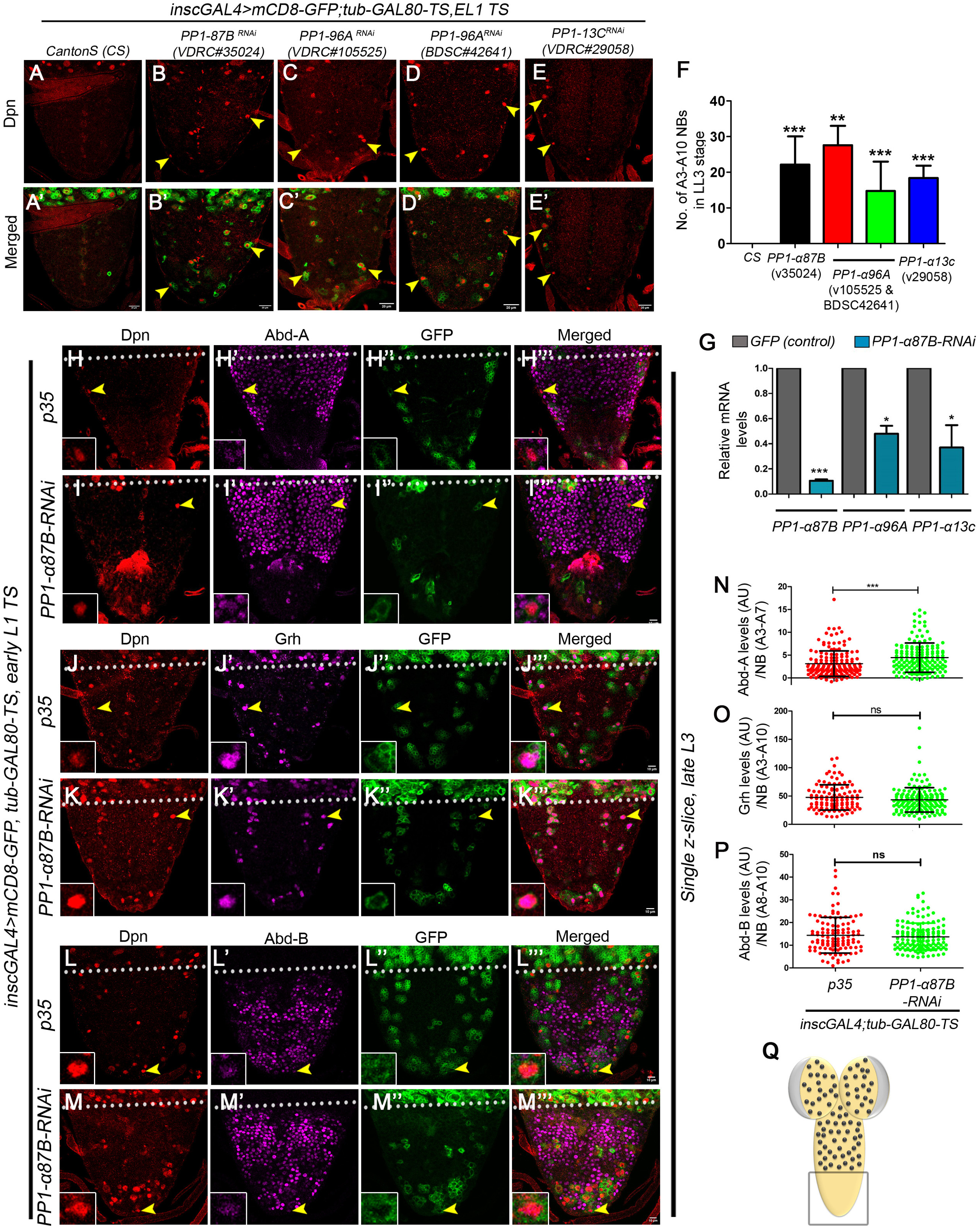
*PP1α* isoforms redundantly regulate NB apoptosis independent of Hox and Grh. (A-E) Shows LL3 stage VNCs of control (*Canton S*) and knockdown for *PP1α* isoforms. Dpn marks the NB, and mCD8-GFP indicates *inscGAL4* expression. At the LL3 stage, control VNCs show no surviving NBs in the A3-A10 segments (A), whereas knockdown of *PP1-α87B* (*VDRC#35024*; 22.17±7.88 NBs, n=6, N=3) (B), *PP1-α96A* (*VDRC#105525, BDSC#342641*; 27.57±5.44 NBs, n=7, N=3 and 14.75±8.24 NBs, n=8, N=3 respectively) (C-D) and *PP1-α13C* (*VDRC#29058;* 18.40±3.43 NBs, n=6, N=3) (E) blocked the NB apoptosis. (F) Show the graph comparing the number of surviving NB in genotypes shown in panels A-E. (G) Shows the graph comparing the relative mRNA levels of different *PP1α* isoforms upon knockdown of *PP1-α87B* (N=3) using the BDSC line (32414). RT-PCR data indicate knockdown of a single isoform (*PP1-α87B*) by RNAi leads to reduction in the RNA levels of other two PP1-α isoforms (*PP1-α13C* and *PP1-α96A*), resulting in an overall decrease of *PP1α*. (H-M) Show that upon the knockdown of *PP1-α87B*, the levels of Abd-B (L-M) and Grh (J-K) are unaffected, while levels of Abd-A (H-I) are increased in the surviving NBs when compared to *p35* expressing controls blocked for NB apoptosis. This establishes that PP1-α87B knockdown blocks NB apoptosis independent of Abd-B, Grh and Abd-A (known triggers of A3-A10 NB apoptosis). In case of Abd-B and Grh no significant change in the levels is observed, in case of Abd-A an increase in levels indicates that the block of death is not due to a reduction in the levels of Abd-A. (N-P) Graph comparing the intensities of Abd-A (*p35*: 3.13±2.79 AU, 157 NBs, n=6, N=3; *PP1-87B-RNAi*: 4.45±3.19 AU, 152 NBs, n=12, N=3) (N), Grh (*p35:* 47.58±22.30 AU for 107 NBs, n=6 VNCs, N=3 vs *PP1-87B-RNAi*: 43.34±21.54 AU for 155 NBs, n=6 VNCs, N=3) (O) and Abd-B (*p35:* 14.41±7.86 AU for 112 NBs, n=7 VNCs, N=3 vs *PP1-87B-RNAi*: 13.74±5.97 AU for 138 NBs, n=7 VNCs, N=3) (P) in A3-A10 NBs in *p35* expressing control VNCs versus VNCs with *PP1-α87B* knockdown. (Q) Region of the CNS shown in all the panels of the figure is indicated by the box. All representative images shown are single confocal sections from LL3-stage female VNCs. Mean intensities are quantified in arbitrary units (AU). Scale bars are 10 µm. Graph shows mean±SD. Significance (P-value) is from One-way ANOVA test and 2-tailed Student’s unpaired t-test. ALH means after larval hatching. Yellow arrowheads indicate NBs. The white dotted line separates thoracic (T) and abdominal (A) segments of the VNC.

**Fig S2:**
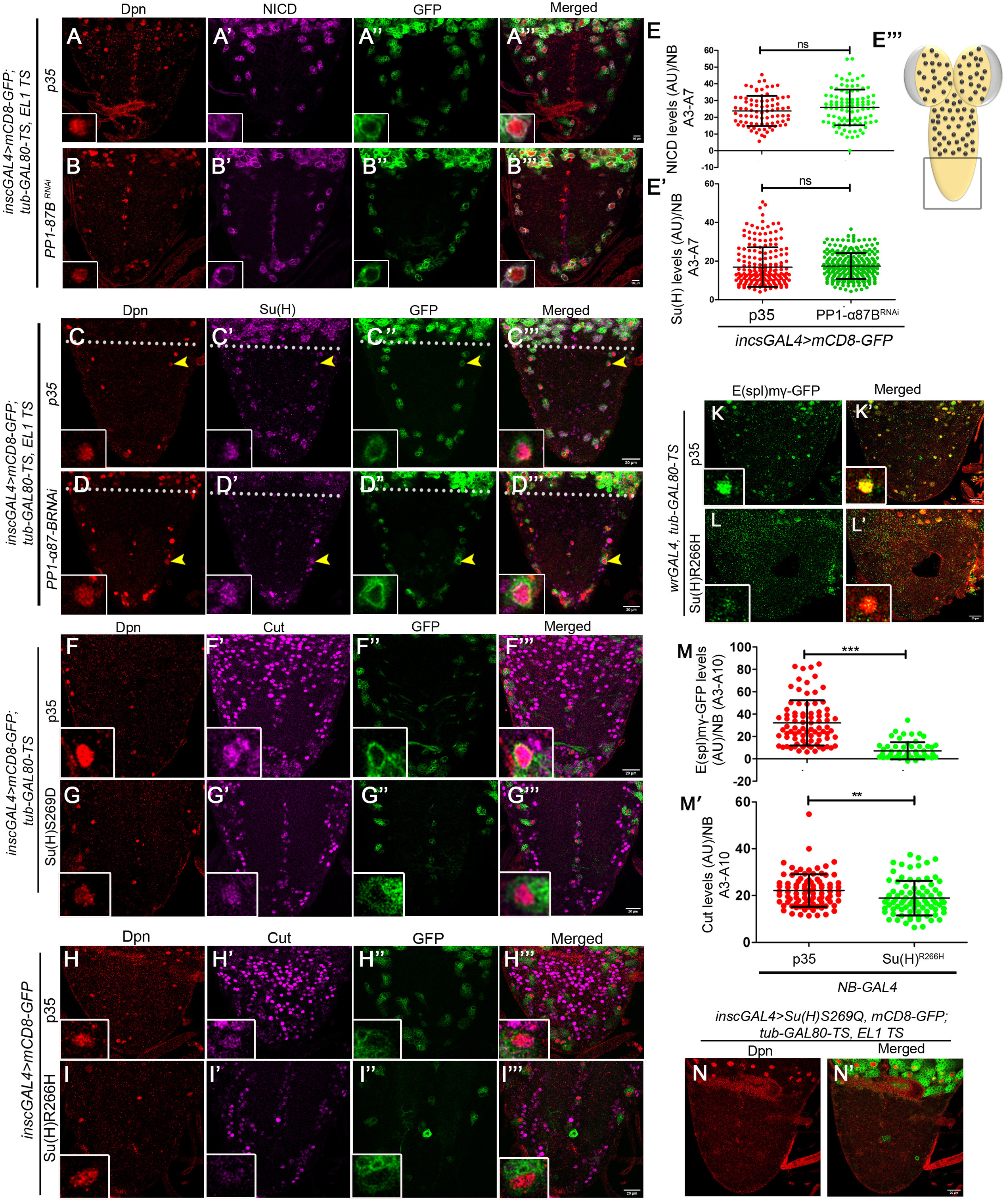
PP1α knockdown does not affect Notch/Su(H) protein levels but alters Su(H) DNA binding capacity to regulate NB apoptosis. (A-B, C-D) Show that upon the knockdown of *PP1-α87B*, the levels of NICD and Su(H), are unaffected in the surviving NBs when compared to p35 expressing controls blocked for NB apoptosis. This establishes that *PP1-α87B* knockdown mediated reduction in the Notch signalling does not affect the levels of NICD and Su(H). (E-E’) Graph comparing the intensities of NICD (*p35*: *23.75±9.01*X±Y AU for 81 NBs, n=6 VNCs, N=3 vs *PP1-α87B-RNAi:* 25.88±10.65 AU for 99 NBs, n=5 VNCs, N=3) (E) and Su(H) (*p35*: 16.90±10.26 for 192 NBs, n=9 VNCs, N=3 vs *PP1-α87B-RNAi*: 17.44±6.76 AU for 253 NBs, n=10 VNCs, N=3) (E’) in A3-A10 NBs in *p35* expressing control VNCs versus VNCs with *PP1-α87B* knockdown. (E’’) Region of the CNS shown in all the panels of the figure is indicated by the box. (F-G, H-I, K-L) Show that upon the overexpression of Su(H)^S269D^ (G) and Su(H)^R266H^ (I and L), the levels of Cut (G and I) and *E(spl)mγ-GFP* (L) are reduced in the surviving NBs in A3-A10 segments, compared to *p35* expressing controls (F, H and K) blocked for NB apoptosis (Cut levels: *inscGAL4>p35*: 50.11±18.49 AU, 149 NBs, n=10, N=3 vs *inscGAL4>Su(H)^S269D^:* 42.19±19.65 AU, 108 NBs, n=13, N=3, graph shown in Fig.1). This establishes that overexpression of Su(H)^S269D^ and Su(H)^R266H^ blocks NB apoptosis by affecting the Notch signalling. (M-M’) Graph comparing the levels of *E(spl)mγ-GFP* (*inscGAL4>p35*: 32.15±20.21 AU, 83 NBs, n=6, N=3; *vs inscGAL4>Su(H)^R266H^*: 7.15±7.65 AU, 60 NBs, n=6, N=3) and Cut levels (*inscGAL4>p35*: 22.16±9.8 AU, 90 NBs, n=8, N=3; *vs inscGAL4>Su(H)^R266H^*: 18.9±7.41 AU, 82 NBs, n=9, N=3) in A3-A10 NBs in *p35* expressing control VNCs versus VNCs with overexpression of Su(H)^R266H^ shown in panels H-I and K-L respectively. (N) Shows that overexpression of Su(H)^S269Q^ (non-phosphorylatable form) does not block the NB apoptosis in A3-A10 segments. All representative images shown are single confocal sections from LL3-stage female VNCs. Mean intensities are quantified in arbitrary units (AU). Scale bars are 20 µm except A-B where it is 10. Graph shows mean±SD. Significance (P-value) is from 2-tailed Student’s unpaired t-test. ALH means after larval hatching. Yellow arrowheads indicate NBs. The white dotted line separates thoracic (T) and abdominal (A) segments of the VNC.

**Fig S3:**
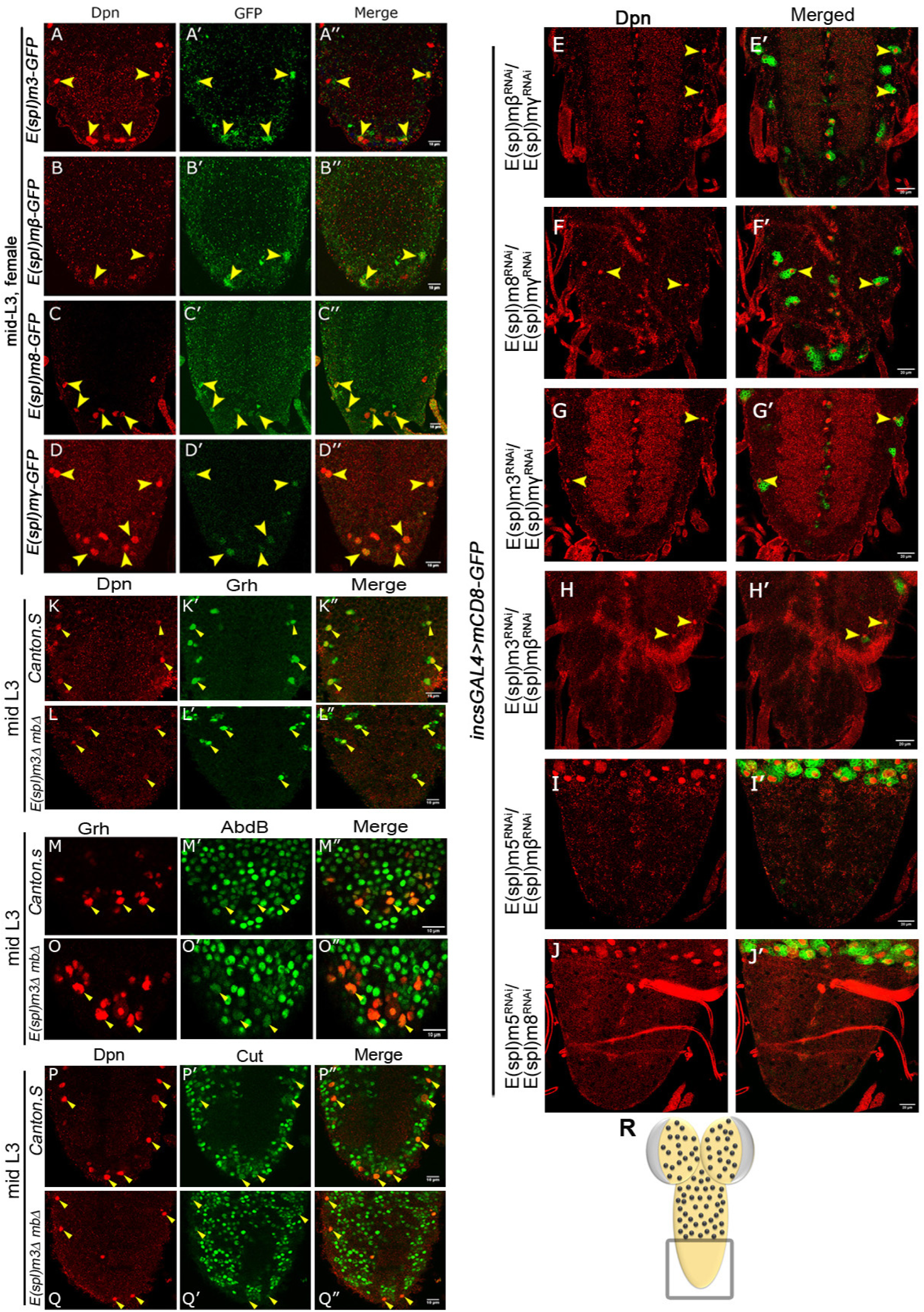
E(spl) complex member proteins are functionally redundant and required for executing NB apoptosis. (A-D) Show the expression of E(spl)-m3, mβ, m8 and mγ protein trap lines in A3-A10 NBs at mid-L3 stage, before their clearance by apoptosis. (E-H) Shows surviving NBs in A3-A10 segments of LL3 stage VNCs for double knockdowns of different members of *E(spl)-complex*: *mβ+mγ* (E), *m8+mγ* (F)*, m3+mγ* (G) and *mβ+m3^i^* (H). (I-J) Show that double knockdowns of *m5+mβ* (I) and *m5+m8* (J) of *E(spl)-complex* do not result in block of NB apoptosis. *E(spl)-m5* is not expressed in A3-A10 NBs, and its knockdown with *mβ* and *m8* does not block NB apoptosis, suggesting that it is dispensable for NB apoptosis. (K-Q) Shows that expression levels of Grh, Abd-B and Cut (Notch target) are unaffected in the NBs in mid-L3 stage VNCs from homozygous double mutant of *E(spl)-Δmß,m3* compared *Canton S* controls (graphs are shown in Fig. 3G-3I). All representative images shown are single confocal sections from mid-and late-L3-stage female VNCs (panels E-J). Scale bars are 10µm for panels A-D, K-Q and 20 µm for panels E-J.

**Fig S5:**
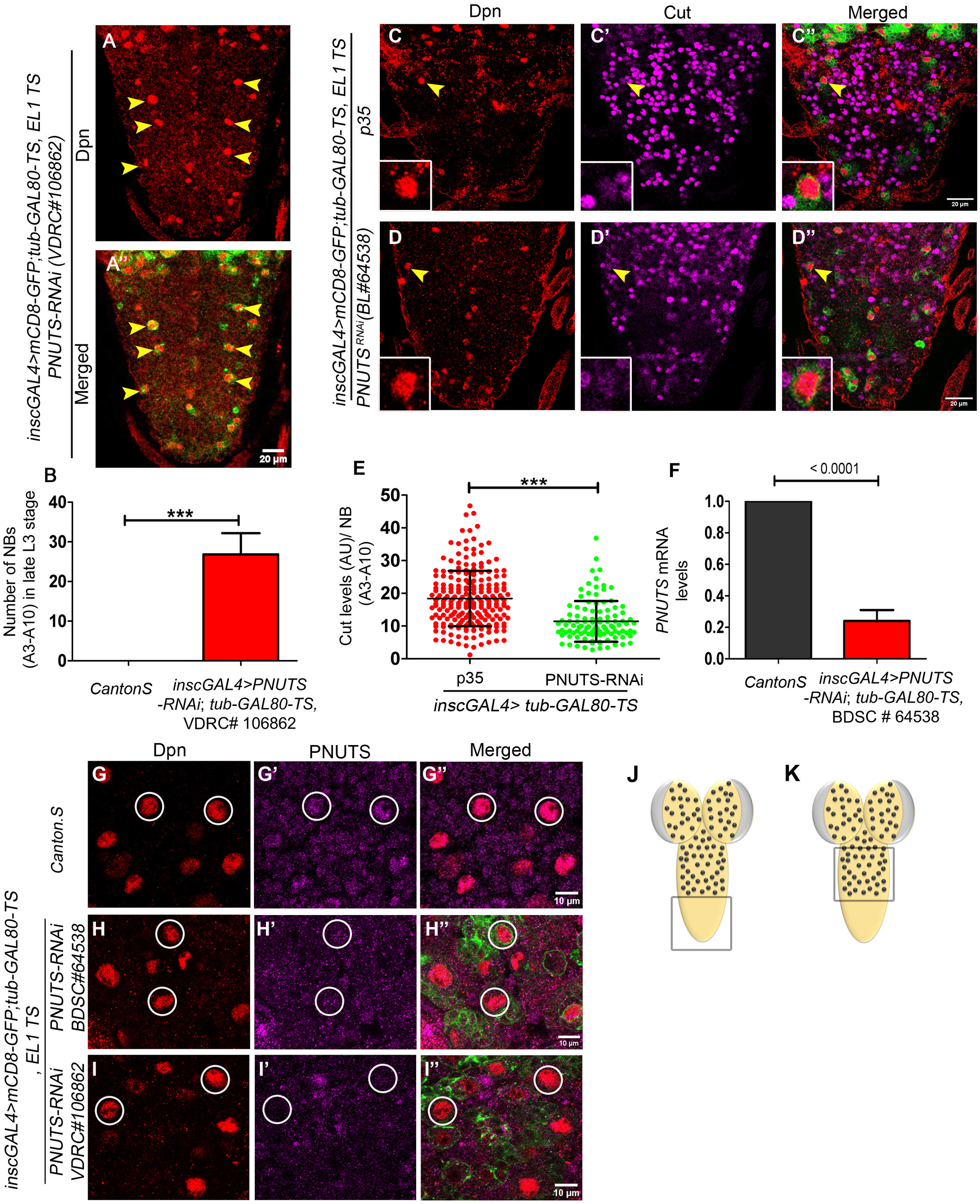
*PNUTS* regulate Neuroblast apoptosis by modulating Notch signalling. (A) Shows the number of surviving NBs (Dpn+ cells) at the LL3 stage in A3-A10 segments of the larval VNC with knockdown of *PNUTS* gene (by VDRC line #106862, same as BDSC line # 64538 shown in Fig. 5). (B) Graph comparing the number of surviving NB in A3-A10 segments for the *Canton S* controls (0 NBs, n=6 VNCs, N=3) vs *PNUTS* knockdown (26.86±5.33 NBs, n=7, N=3). (C-D) Show that upon the knockdown of *PNUTS*, the levels of Cut are reduced in the surviving NBs when compared to *p35*-expressing controls blocked for NB apoptosis. Insets show magnified view of the NBs indicated by arrowheads. This suggests that *PNUTS* knockdown by VDRC line also blocks NB apoptosis by affecting Notch signalling. (E) Graph comparing the intensities of Cut levels in A3-A10 NBs in p35 (18.39±8.45 AU, 200 NBs, n=8, N=3) expressing control VNCs versus VNCs with *PNUTS* knockdown by BDSC # 64538 (11.42±6.22 AU, 106 NBs, n=9, N=3). (F) Shows the graph comparing the relative mRNA levels of *PNUTS* mRNA upon knockdown of the *PNUTS gene by BDSC line #64538* (N=3) used in all the experiments. (G-I) Compares the PNUTS antibody staining, in the larval thoracic NBs in case of control (*inscGAL4>Canton.S*) (G), and the knockdown for *PNUTS* (*inscGAL4>PNUTS*-*RNAi)* by two independent RNAi lines: BDSC line (#64358) (H), and VDRC line (#106862) (I); establishing that knockdown results in a reduction in PNUTS levels in thoracic NBs. (J-K) Regions of the CNS shown in figure panels A, C-D (J) and panels G-I (K) are indicated by the boxes. All representative images shown are single confocal sections from LL3-stage female VNCs. Mean intensities are quantified in arbitrary units (AU). Scale bars are 20 µm for panels A, C-D and 10 μm for G-I. Graph shows mean±SD. Significance (P-value) is from 2-tailed Student’s unpaired t-test. Yellow arrowheads and white circles indicate NBs. The white dotted line separates thoracic (T) and abdominal (A) segments of the VNC.

**Fig S6:**
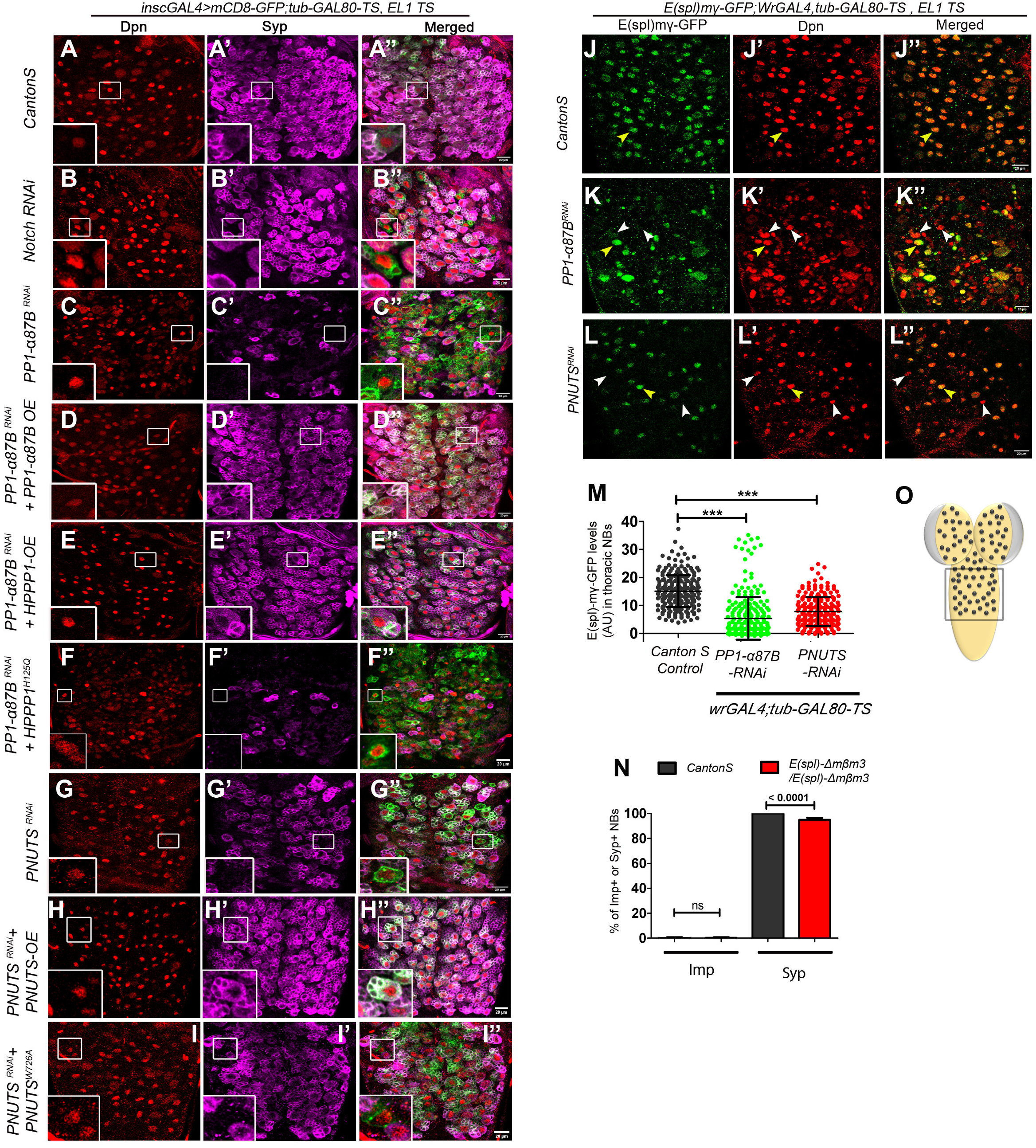
PP1-α/PNUTS is required for the NBs transition to the late factor Syncrip. (A-I) Shows the thoracic segment of LL3 stage larvae of the indicated genotypes. Brains are immunostained with late temporal factor Syncrip (Syp shown in Magenta) and NB marker Dpn (Red). (A) All the NBs in the control (Canton S) VNCs express Syp by the LL3 stage (100% NBs, n=9 VNCs, N=3), but upon *Notch* knockdown (B), 11% of thoracic NBs fail to express Syp, indicating their inability to undergo the competence switch, remaining NBs express Syp (89.41±4.47% NBs Syp+, n=13, N=3) (Fig. 1J). (C) Knockdown of *PP1-α87B* results in 50% of NBs failing to express Syp at the LL3 stage, while the remaining 50% NBs express Syp normally (50.63±6.78% NBs; n=8 VNCs; N=3. This defect is rescued by overexpression of *Drosophila* (*PP1-α87B*: 97.87±2.571% NBs Syp+; n=7 VNCs; N=3) (D), and human ortholog (hPPP1: 100±0% NBs Syp+, n=7 VNCs, N=3) of PP1α (E), but not by phosphatase dead version of the human ortholog (hPPP1^H125Q^: 44.50±8.96% NBs Syp+, n=10, N=4), establishing the specificity and suggesting a cross phylum conservation of the phenotype. (G) Shows that 13% of thoracic NBs (n=9 VNCs, N=3) fail to express Syp, indicating their inability to undergo the competence switch, while the remaining 87% NBs express Syp (87.82±1.19% NBs Syp+, n=12, N=4). (H-I) This defect is rescued by overexpression of wild-type *Drosophila PNUTS*, where all the NBs expressed Syp (H), but not by the PP1-α-binding-deficient form of *PNUTS* (*PNUTS^W726A^*: 83.88±4.29% NBs Syp+, n=13VNCs, N=3), where only 83% NBs expressed Syp, while the rest 17% NBs did not express Syp, establishing the specificity of the phenotype and suggesting that PNUTS need to interact with PP1α to rescue the phenotype. (J-L) Show that compared to controls (*Canton S*) (J), upon the knockdown of *PP1-α87B* (K) and *PNUTS* (L), the levels of *E(spl)mγ-GFP* are reduced in thoracic NBs. This suggests that *PP1-α87B* and *PNUTS* knockdown affecting Notch signalling in thoracic NBs. (M) Graph comparing the intensities of *E(spl)mγ-GFP* in thoracic NBs in Canton S VNCs (15.10±5.69 AU, 248 NBs, n=10, N=3) versus VNCs with *PP1-α87B* (5.36±7.60 AU, 253 NBs, n=6, N=3) and *PNUTS* (7.82±5.15 AU, 224 NBs, n=8, N=3) knockdown. (N) Graph showing the percentage of thoracic NBs expressing early factor Imp and late factor Syp in the case of control VNCs (*CantonS)* and homozygous test VNCs (*Δmβm3/Δmßm3*). The *E(spl)-Δmß,m3* double deletion does not affect E>L transition. (O) Region of the CNS shown in all the panels of the figure is indicated by the box. All representative images shown are single confocal sections from thoracic region of LL3-stage female VNCs. Mean intensities are quantified in arbitrary units (AU). Scale bars are 20 µm. Graph shows mean±SD. Significance (P-value) is from one way ANOVA test. ALH means after larval hatching. The insets in panels A-I are the magnified view of the NBs indicated by white boxes.

**Fig. S7.**
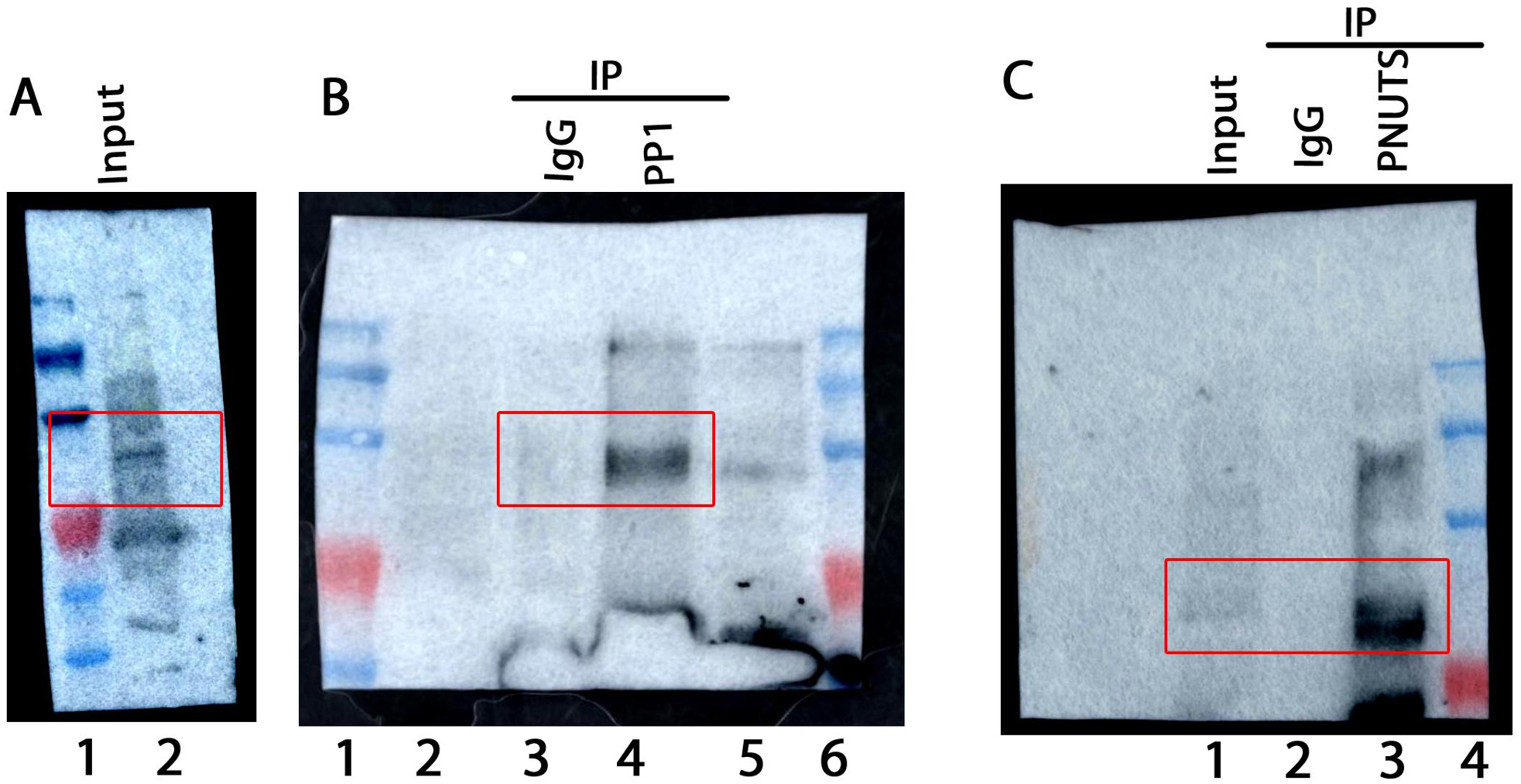
PP1 and PNUTS interacts with Su(H) *in vivo*. Uncropped original pictures of the western blot showing that PP1 could pull down Su(H). (A) Lane 2 is input probed with anti-Su(H) antibody. (B) Lane 3 is mock IgG and lane 4 is PP1 IP. The blots are probed with anti-Su(H) antibody. Uncropped original pictures of the western blot showing that PNUTS could pull down Su(H)(A). Lane 1, 2 and 3 is lysate input, mock IgG and PP1 IP respectively. The blot is probed with anti-Su(H) antibody.

## Acknowledgements

We thank F. Schweisguth, Anette Preiss, Anja C Nagel, J. Knoblich, Girish Ratnaparkhi for reagents; CDFD animal facility, Bioklone Biotech Pvt. Ltd., Chennai and the Developmental Studies Hybridoma Bank (DSHB) at The University of Iowa for antibodies, Sophisticated Equipment Facility at CDFD for DNA sequencing and TFF at NCBS, Bangalore for transgenic flies. We thank M.S. Reddy and S. Tyagi for suggestions, A. Kunchur for generating Imp and Syp antibodies, C. S. Singh and Feroze Syed for their assistance in various phases of the project, as well as the past and current member of LNCB for their comments on the work during group meetings. This study was funded by; Department of Science and Technology, India (CRG/2021/003275); Department of Biotechnology, India (BT/PR41306/MED/122/ 259/2020; BT/PR45460/MED/12/952/2022, BT/54952/BMS/85/365/2024); CDFD core funds; and UGC-JRF (award to JB) [UGC-No: F.16-6(DEC.2016)/2017(NET)], UGC SRF Award to A.B. [UGC Ref No. 22/06/2014(i)EU-V, 2061430472] and ICMR, India (award to R.S.) [ICMR Ref.No:3/1/3/JRF-2012/HRD-63 (40260)].

## Author contribution

RJ and JB conceptualised the study; RS made the initial observation and preliminary experiment; JB did most of the experiments; AB contributed to E(spl) deletion data; JB, AB, RS and RJ analysed the data; JB, AB made the figures; and RJ and JB wrote the manuscript.

## Supplementary Materials and Methods

### Fly stocks

The following additional fly lines were used: *PP1-a87B* RNAi (VDRC-35024, BDSC-67911), *PP1-a96A* RNAi (VDRC-105525, BDSC-42641), *PP1-a13C* RNAi (VDRC-29058), *E(spl)my-GFP E(spl)m3-GFP, E(spl)m{3-GFP, E(spl)m8-GFP* {Couturier, 2019 #390}.

### Molecular cloning

*PP1-a87B, Imp,* and *Syncrip f*ull-length cDNAs were gifts from the Ratnaparkhi lab (@llSER Pune, amplified from DGRC clones LD03380, BS07088, and BS07872) and sub-cloned into the Ndel and Xhol sites of the pETM11 vector (modified from pET14b). Full-length *PNUTS* was generated by performing overlap extension PCR from the Expressed-sequence tags (EST) plasmids of *PNUTS* gene. Briefly, the N-terminal (1-1708 bp) and C-terminal (1682-3408 bp) fragments of *PNUTS* with 27 bp common sequence were PCR amplified using Expressed-sequence tags (EST clones from DGRC-RH48883 and LD47649). Subsequently, overlap extension PCR was done to obtain a full-length coding sequence of *PNUTS* and subsequently cloned in pCR 2.1-TOPO TA vector (lnvitrogen)*. Su(H)* wildtype and different mutant constructs were PCR amplified and cloned in pUASTattB-HA plasmid using lnFusion HD cloning kit (Takara Clontech). *Su(H)^WT^* and *Su(H)^S269D^* and *Su(H)^R266H^* were PCR amplified using genomic DNA from transgenic flies gifted by Anja C Nagel {Frankenreiter, 2021 #383}. A similar strategy was also applied to clone *PP1-a87B* (RNAi resistant for BSDC-32414), *PNUTS^WT^* (RNAi resistant for BDSC-64538)*, PNUTS^W726A^* (RNAi resistant for BDSC-64538), hPPP1^H1250^ in pUASTattB-HA plasmid. Primer list: **Table 2**.

**Table 2:**
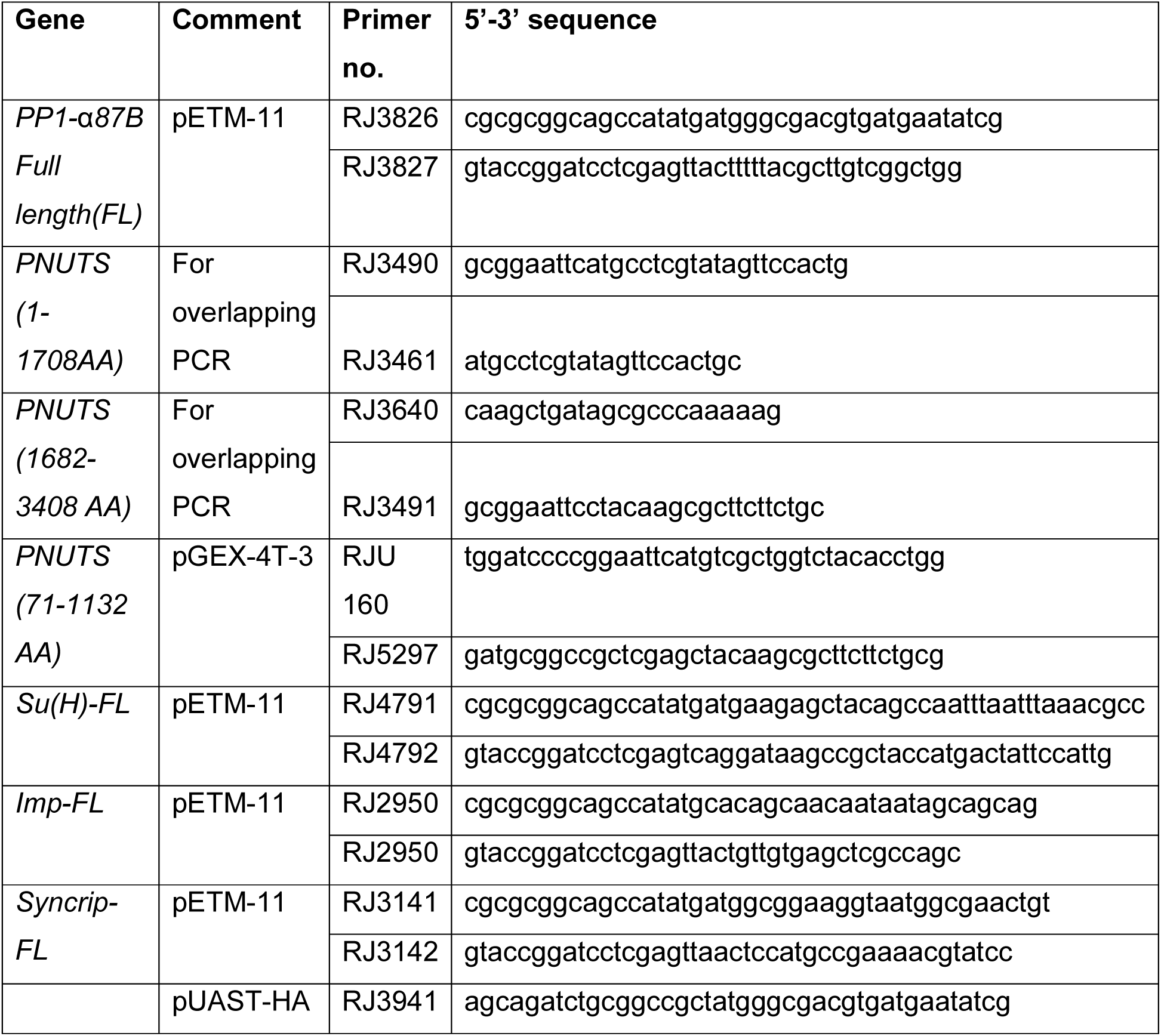
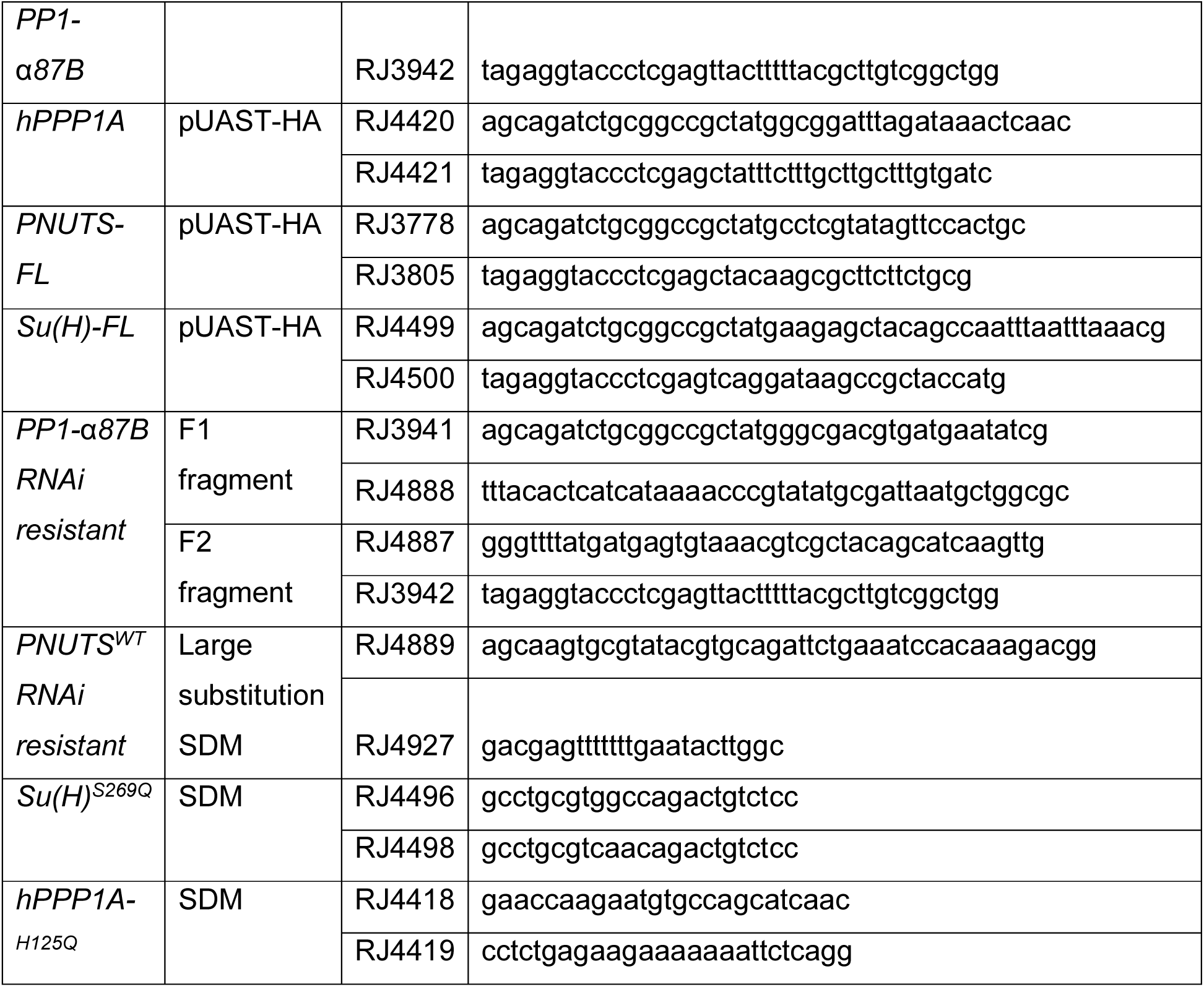
List of primers used for SDM and subcloning.

### Site directed mutagenesis

To generate *PNUTS^W726A^, Su(H)^S269Q^* and *hPPP1^H125Q^* constructs, non-overlapping primer-based site-directed mutagenesis PCR was used. Briefly, a pair of non-overlapping primers was used to amplify the entire circular plasmid template and the forward primer carrying the desired mutation. Further, the PCR product was digested with the Dpnl enzyme to remove the parental template, followed by the ligation and transformation. Primer list: **Table 2**.

### Generation of RNAi-resistant *PP1-a87B* and *PNUTS* construct

The *PP1-a87B* and *PNUTS* nucleotide sequences targeted by RNAi fly lines: BDSC-32414 and BDSC-64538 respectively, were altered without changing the protein sequence (primers given in Table 2). To generate the RNAi-resistant construct of *PP1-a87B* two-template overlap extension PCR strategy was utilized. Briefly, the N-terminal and C-terminal fragments of *PP1-a87B* with a modified RNAi-targeted region as a common overlap were PCR amplified. Subsequently, overlap extension PCR was done using these two PCR products as templates to obtain a full-length RNAi-resistant coding sequence of *PP1-a87B*. To generate the RNAi-resistant construct of *PNUTS*, a non-overlapping primer-based SDM PCR strategy was utilized, with the forward primer (detailed in Table 2) carrying the modified RNAi-targeted region as 5’ overhang.

### Antibody generation

To generate anti-PP1-a87B, PNUTS, lmp and Syncrip antibody, the full length coding sequence of *PP1-a87B,Imp, Syncrip* and 1-879 bp of *PNUTS* with His-Tag at the N-terminus respectively was cloned in *pETM-11* plasmid (modified from pET14b) using lnFusion HD cloning kit (Takara Clontech) and further expressed in BL21 *E. coli* cells. The purified proteins were used to immunize mice using standard protocols to generate polyclonal antisera.

### Generation of transgenic flies

The following HA tagged constructs cloned in pUASTattB-HA plasmid were inserted in a site-specific manner either at attP40-25C6 (Chr 2) or attP2-68A4 (Chr 3) position: *UAS-Su(H)^WT^*, *UAS-Su(H)^S269D^*, *UAS-Su(H),UAS-Su(H)^R266H^*, *UAS-PP1-a87B*(RNAi resistant for BSDC-32414), *UAS-PNUTS^WT^*(RNAi resistant)*, UAS-PNUTS^W726A^* (RNAi resistant for BDSC-64538),*UAS-hPPP1^H125Q^*.

### RNA isolation and qRT-PCR

Total RNA was extracted using 40-45 L3 larval brains of appropriate genotypes. TRlzol (Thermo Fisher #15596018), followed by chloroform extraction and isopropanol precipitation was used to extract the RNA. To remove the residual genomic DNA, RNA was further treated with DNase (lnvitrogen TURBO DNA-free kit AM1907) and then cDNA was synthesised from 0.5µg of RNA using Takara cDNA synthesis kit (PrimeScript cDNA Synthesis Kit #6110A). cDNA was diluted 10-fold before using it for qRT-PCR. cDNA was then subjected to qRT-PCR using Takara SYBR Green (TB Green Premix Ex Taq, ROX Plus #RR42LR) in Himedia LA1074-1NO lnsta 096 - 6.0. Transcript levels were quantified using 2^-LiLiCt^ method, and transcript levels were normalized using *GAPDH* housekeeping gene. Primer sequences are provided in the supplementary data **Table 3.**

**Table 3:**
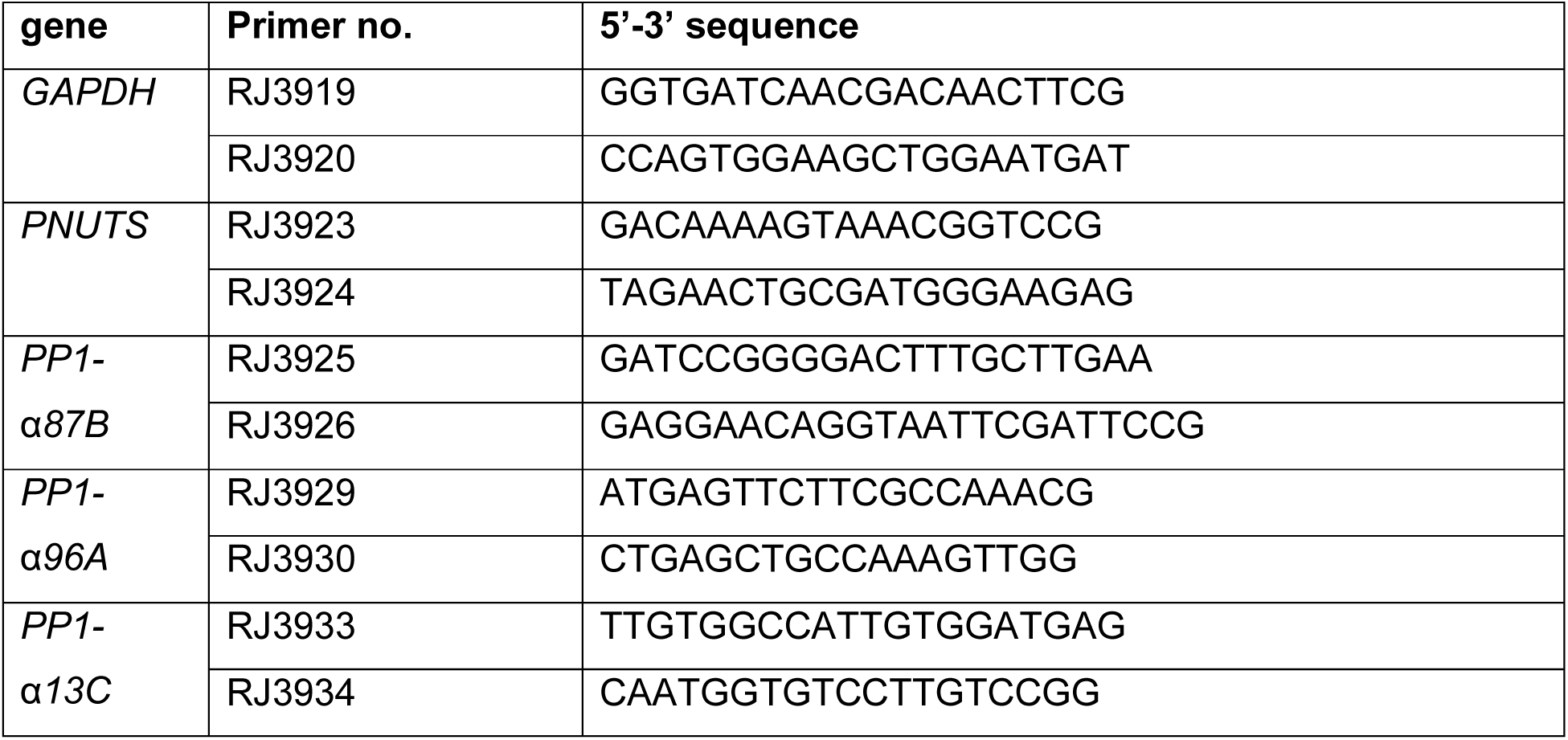
List of primers used in RT-PCR.

